# The ESCRT-III proteins IST1 and CHMP1B assemble around nucleic acids

**DOI:** 10.1101/386532

**Authors:** Nathaniel Talledge, John McCullough, Dawn Wenzel, Henry C. Nguyen, Matthew S. Lalonde, Monika Bajorek, Jack Skalicky, Adam Frost, Wesley I. Sundqust

## Abstract

ESCRT-III proteins can promote inside-out or outside-in membrane tubulation and fission. In addition, several observations suggest that ESCRT factors may also associate with nucleic acids during development, different stages of the cell cycle, and during retro-transposition of parasitic nucleic acids like LINE1 elements. Two ESCRT-III subunits, IST1 (aka CHMP8) and CHMP1B, can coassemble as an external protein coat around liposomes *in vitro* and around recycling endosomal tubules in living cells. Here we show that recombinant IST1 and CHMP1B can also copolymerize into double stranded filaments that surround nucleic acids. Electron cryo-microscopy reconstructions of nucleic acid-bound IST1-CHMP1B copolymers revealed that the polynucleotides track along a binding groove formed between filaments of the inner CHMP1B strand. The well-ordered structures also reveal that the C-terminal tails of CHMP1B subunits extrude through the outer IST1 layer to the tube exterior. As a result, the MIT domain binding motifs of both CHMP1B and IST1 are arrayed on the outer surface of the copolymer, where they could bind and recruit MIT domain-containing co-factors, such as the SPASTIN ATPase or the USP8 ubiquitin protease. Our structure raises the possibility that ESCRT-III proteins may form nucleic acid complexes in mammalian cells.

## INTRODUCTION

The endosomal sorting complexes required for transport (ESCRT) pathway functions in diverse membrane remodeling processes (Campsteijn et al., 2016; Christ et al., 2017; Frankel and Audhya, 2017; Lippincott-Schwartz et al., 2017; McCullough et al., 2018; Schoneberg et al., 2017; Scourfield and Martin-Serrano, 2017; Stoten and Carlton, 2018). The ' early-acting ‘ ESCRT factors ALIX (ALG2-interacting protein X) and the ESCRT-I/II complexes, link cargo recognition to recruitment and activation of 'late-acting’ ESCRTs, which include filament-forming ESCRT-III complexes and ATPases associated with diverse cellular activities (AAA ATPases), like VPS4 (vacuolar protein sorting 4) and SPASTIN. Recruitment to the target membrane by early-acting components activates ESCRT-III proteins to polymerize into membrane-bound filaments. These filaments cooperate with the ATPases to constrict the membranes and catalyze membrane fission. ESCRT-dependent membrane fission events occur in a still growing list of cellular processes, including the well-characterized pathways of multivesicular body (MVB) formation (Bryant and Stevens, 1998; Henne et al., 2013), cytokinetic abscission (Scourfield and Martin-Serrano, 2017; Stoten and Carlton, 2018), and HIV and other virus egress pathways (Votteler and Sundquist, 2013). In such “inside-out” membrane fission reactions, the membrane neck encloses cytoplasm and the ESCRT-III filaments assemble within the membrane neck, where they draw opposing membranes together to drive membrane fission.

Recent work indicates that at least a subset of the 12 human ESCRT-III proteins can also catalyze membrane-shaping reactions of the opposite orientation: driving "outside-in" membrane fission reactions, rather than the cannonical ESCRT-dependent "inside-out" reactions described above. Specifically, the ESCRT-III proteins CHMP1B and IST1/CHMP8 (increased sodium tolerance 1, aka CHMP8) can co-polymerize into double-stranded filaments that can stabilize positive membrane curvature *in vitro* and co-localize around the cytoplasmic surface of endosomal tubules within cells (McCullough et al., 2015). *In vitro* CHMP1B binds to and remodels liposomes into tapering tubes by wrapping around their exterior. In cells, CHMP1B and IST1 filaments can coat membrane tubules that extend into the cytoplasm (McCullough et al., 2015). Vesicles that recycle endosomal proteins back to the plasma membrane can be released from SNX1-positive endosomal tubules that contain IST1 and SPASTIN (Allison et al., 2017; Allison et al., 2013). These observations support a model in which IST1-CHMP1B cofilaments can work together with the AAA ATPase SPASTIN to constrict endosomal membranes toward the fission point (Allison et al., 2017; Allison et al., 2013; McCullough et al., 2015), although this model remains to be tested. Recent work suggests that the ESCRT pathway may also play an analogous role in budding nascent peroxisomal vesicles from ER membranes, although this process is not yet well understood (Mast et al., 2018).

Despite considerable sequence diversity, all 12 different human ESCRT-III proteins appear to contain a structurally similar core (comprising helices α1-α5) (Bajorek et al., 2009b; Muziol et al., 2006; Xiao et al., 2009). In at least some ESCRT-III subunits, this core region can interconvert between a soluble monomeric conformation (the “closed” conformation), and an assembled membrane-bound conformation (the “open” conformation) (Lata et al., 2008; Lin et al., 2005; McCullough et al., 2015; McMillan et al., 2016; Tang et al., 2015). Both protein-protein interactions and post-translational modifications, including ubiquitination, appear to regulate interconversion of the two conformational states. For example, filament formation is typically nucleated by interactions with upstream ESCRT factors (Christ et al., 2017; Schoneberg et al., 2017), whereas ubiquitination of CHMP1B has been reported to prevent CHMP1B from opening or polymerizing (Crespo-Yanez et al., 2018).

Residues beyond the ESCRT-III core typically lack persistent structure, but contain binding sites for interactions with both upstream (BRO domain proteins) and downstream (microtubule interacting and transport (MIT)-domain containing proteins) binding partners. The sequence elements responsible for MIT domain binding are known as ‘MIT interacting motifs’, or MIMs. A series of crystal and NMR structures have revealed a remarkable diversity of different MIT-MIM binding modes, suggesting that these interactions may comprise a recognition "code" that dictates which ESCRT-III subunits can recruit which MIT domain cofactors (Caballe et al., 2015; Kieffer et al., 2008; Obita et al., 2007; Skalicky et al., 2012; Solomons et al., 2011; Stuchell-Brereton et al., 2007; Yang et al., 2008).

A striking feature of the IST1-CHMP1B copolymer is the highly basic nature of its lumenal surface, which is created by a series of conserved lysine and arginine residues located on CHMP1B helix 1. This helix was proposed to be the major membrane-binding surface in other ESCRT-III proteins (Buchkovich et al., 2013; Muziol et al., 2006), and we found that this helix of CHMP1B binds to negatively charged membrane surfaces (McCullough et al., 2015).

We sought to test whether the electropositive interior of the IST1-CHMP1B copolymer can bind other polyanions such as nucleic acids. In part, these studies were motivated by the intriguing history of reported associations between ESCRT proteins and nucleic acids. Potentially relevant observations include: 1) The original description of the ESCRT-III protein CHMP1A reported that it localized to chromatin, and appeared to play a role in creating regions of nuclease-resistant, condensed chromatin and associated with the transcriptional repressor polycomb-like protein (Pcl) on condensed chromatin (Stauffer et al., 2001). 2) Both CHMP1A and CHMP1B contain nuclear localization signals (Manohar et al., 2011; Yang et al., 2004). 3) CHMP4B localizes to chromosome bridges and micronuclei, and co-immunoprecipitates with chromatin (Sagona et al., 2014). 4) ESCRT deletions can cause defects in chromosomal segregation in humans and yeasts (Reid et al., 2011). 5) CHMP2A (also known as breast adenocarcinoma 2 (BC-2)) was identified in the nuclear matrix, and its overexpression induces condensed chromatin (Hodges et al., 2005). 6) The Bioplex network database reports interactions between CHMP1A and a number of homeobox-containing transcription factors (Schweppe et al., 2018). 7) Finally, long interspersed nuclear element-1s (LINE-1s, or L1s), the active retrotransposons and mutagens of mammalian genomes, depend on poorly understood interactions with ESCRTs, including ESCRT-III proteins (Horn et al., 2017). ESCRT-I and -II factors have also been reported to interact with nucleic acids (Emerman and Blower, 2018; Irion and St Johnston, 2007; Moriscot et al., 2011).

Here, we report that nucleic acid polymers are potent nucleators of IST1-CHMP1B filament formation. The resulting protein coat enwraps the nucleic acids and protects them from nucleases. High-resolution cryoEM reconstructions of the nucleic acid-stimulated structure demonstrated that the structure matches that of the nucleic acid-free CHMP1B-IST1 polymer (McCullough et al., 2015), and additionally revealed that the C-terminal MIM1 helix of CHMP1B plays a key role in copolymerization. A flexible linker between the domain-swapped helix α5 of CHMP1B enables the sixth helix, which contains the MIM1 element, to extend out to the exterior surface of the coassembly, where it packs against the closed-conformation fifth alpha helix of IST1. Snorkeling of the CHMP1B terminus to the outer surface also makes the MIM helix available to other ESCRT-III binding partners, including MIT domain proteins like the AAA ATPase SPASTIN and the ubiquitin-specific protease USP8, both of which recognize the MIM1 tail of CHMP1B and appear to regulate its assembly status.

## RESULTS

### Nucleic acids promote assembly of an IST1-CHMP1B copolymer

We have previously shown how recombinant IST1 and CHMP1B can coassemble into helical tubes comprising double-stranded filaments, both free and on negatively-charged membrane tubules (McCullough et al., 2015). To test whether nucleic acids can also bind and nucleate the assembly of these ESCRT-III proteins, we identified physiological solution conditions (125 mM NaCl, 25 mM Tris pH 8.0, 4 μM [IST1] and 4 μM [CHMP1B]) that do not support coassembly of IST1 and CHMP1B owing to the low protein concentration and elevated salt concentration (**Figure 1A**). We then tested whether addition of linear dsDNA, ssDNA, and/or ssRNA could nucleate ESCRT-III polymerization under these solution conditions. As shown in **Fig. 1B-D**, we observed that all three types of nucleic acid potently stimulated IST1-CHMP1B copolymer formation.

**Figure 1:**
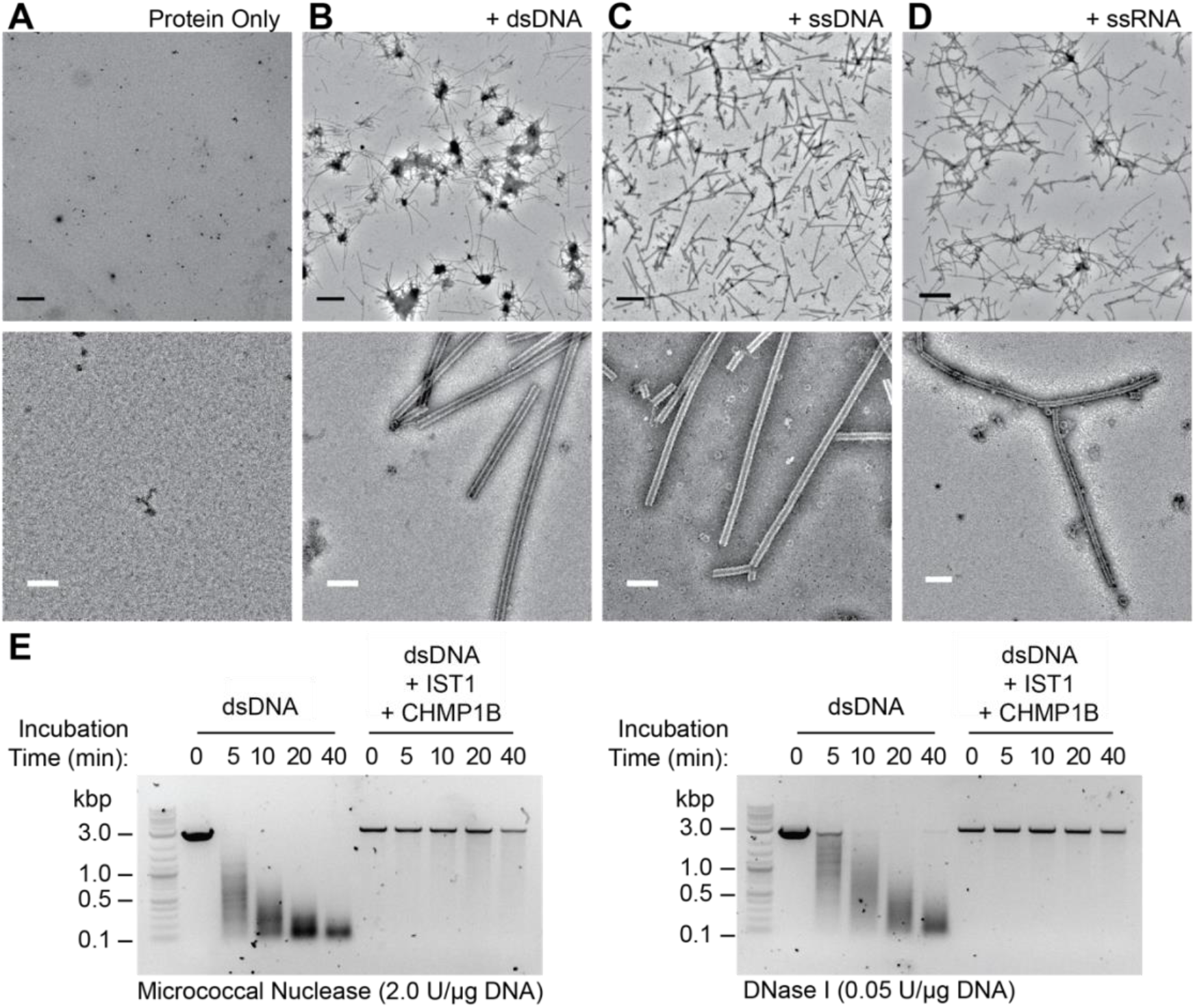
Nucleic acids are encapsulated within CHMP1B and IST1 copolymers. (**A-D**) Negatively stained electron micrographs of IST1-CHMP1 B copolymer mixtures incubated under physiological salt conditions (125 mM NaCl), either alone (**A**) or with dsDNA (**B**), ssDNA (**C**) or ssRNA (**D**). Low (upper row) and medium (bottom row) magnification images are shown with 1 μm scale bars (black) and100 nm scale bars (white). (**E**) dsDNA samples incubated the absence or presence of IST1_ntd_-CHMP1B, treated with the designated nucleases, and imaged following agarose gel electrophoresis.

We next tested whether the nucleic acid-bound copolymers were merely nucleating a protein-only assembly or whether the protein bound and wrapped the nucleic acid polymer, much as it would an anionic membrane. To answer this question, we performed nuclease protection assays to compare the accessibility of dsDNA that was either free (control condition) or in complex with the IST1-CHMP1B copolymer. As shown in Fig. 1E, free DNA was readily degraded by both micrococcal nuclease and DNase I, so that no full-length polynucleotides were detectable after 40 minutes. In contrast, full-length dsDNA was completely protected by the IST1-CHMP1B copolymer, even after 40 minutes. Hence, dsDNA, IST1 and CHMP1B copolymerize together to form a structure in which the DNA is protected from nuclease digestion, likely because it resides within the lumen of the helical protein tube.

To verify that the nucleic acids were protected within the tube lumen, we used cryoEM to visualize all three of the nucleic acid bound IST1-CHMP1B complexes. In all cases, additional density corresponding to nucleic acid was visible within the lumen of the copolymer filaments (**Figure 1-figure supplement 1A**). Furthermore, two-dimensional classification of overlapping particle segments of the ssDNA-bound cofilaments revealed that the DNA density lined the ‘grooves’ between the CHMP1B strands within the interior of the copolymeric coat (**Figure 1-figure supplement 1B**).

### High-resolution reconstruction of the ssDNA-bound IST1-CHMP1B copolymer

To visualize precisely how nucleic acids bind within the helical IST1-CHMP1B filaments, we determined the 3D structure of the ssDNA-IST1-CHMP1B complex using cryoEM and helical reconstruction (**Table 1**). This sample utilized full-length, human IST1 (1–366) and a nearly full-length human CHMP1B construct (4–199), incubated with a single-stranded poly(dAdG)_200_ oligonucleotide (**Figure 2-figure supplement 1 A-B**) (McCullough et al., 2015). This sample was well-ordered and the overall resolution of the protein-only portion of the 3D reconstruction reached 2.9 Å, as judged by gold-standard FSC criteria (Materials and Methods and **Figure 2-figure supplement 2**). The overall structure of the protein helical assembly was similar to our previously published structure of the protein-only assembly of IST1-CHMP1B (McCullough et al., 2015), showing a double-stranded protein filament with a helical rise of ~3.2 Å and a twist of ~21.2°. Although the new assemblies contain full-length IST1 _1–366_, (rather than the the N-terminal ESCRT-III domain composed of residues 1–189 (IST1_NTD_), used previously), no additional density was observed beyond the IST1 ESCRT-III core helices (α1- α5). Thus, the C-terminal half of IST1 from residue ~180 to 366 remains unstructured and has a high degree of flexibility, even when present in the assembled copolymer.

**Figure 2:**
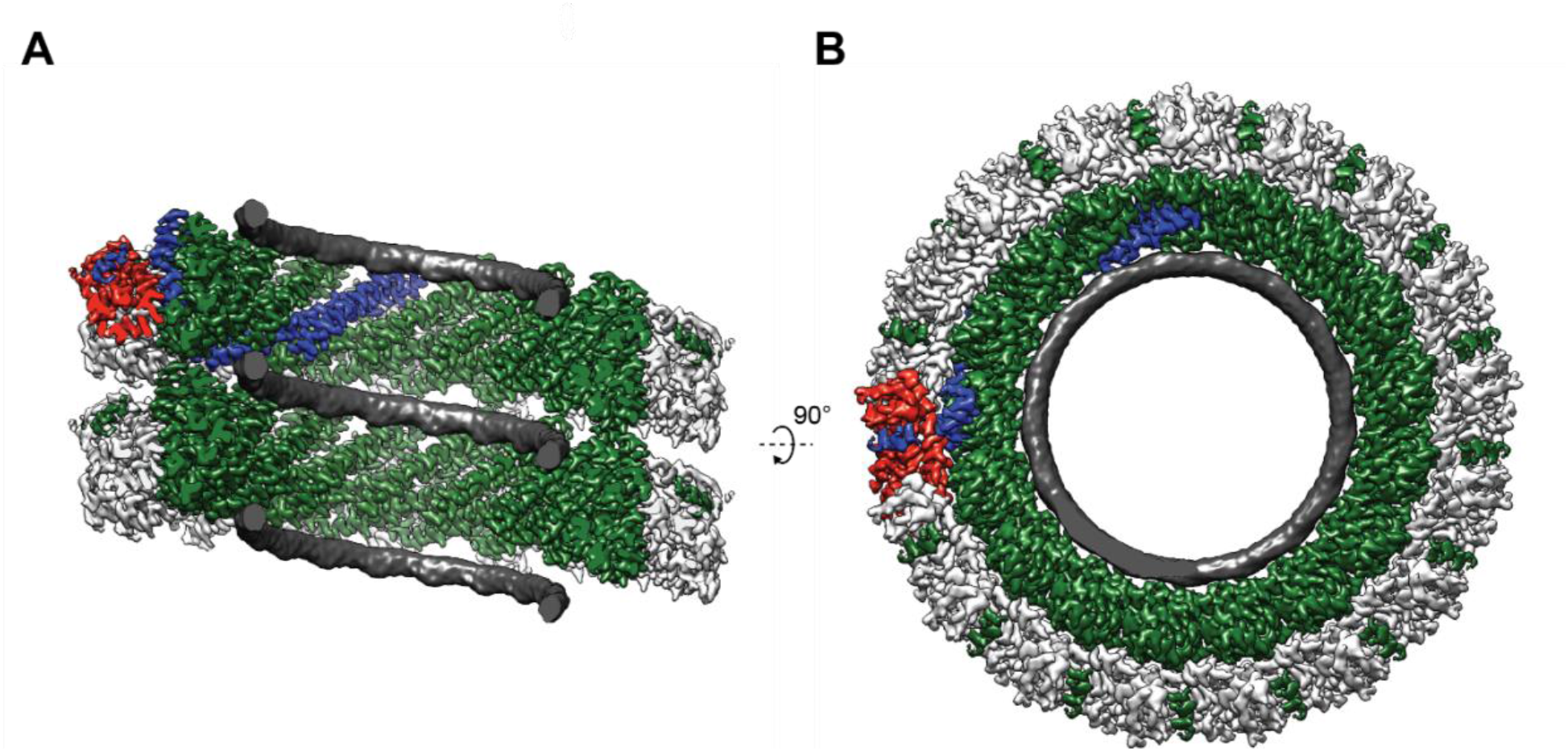
CryoEM density of ssDNA bound within a copolymer of full-length IST1 and CHMP1B. (**A**) Internal cutaway view of the reconstructed heteropolymeric double-stranded helical assembly of IST1 (white, outer strand) and CHMP1 B (dark green, inner strand) encapsulating ssDNA (dark grey). (**B**) End-on view looking down the helical axis of the reconstruction. A single CHMP1B (blue) and an IST1 (red) subunit are highlighted in both views of the reconstruction.

**Table 1:** Overview of the CryoEM Data Collection and Validation Parameters.

| Microscopy |  |  |  |
| --- | --- | --- | --- |
| Dataset | ssDNA<br>EMD-9005 | dsDNA | ssRNA<br>(Data not shown) |
| Microscope | Titan Krios | TF20 | TF20 |
| Voltage (kV) | 300 | 200 | 200 |
| C2 aperture ($\mu\text{m}$) | 40 | 70 | 70 |
| Objective aperture ($\mu\text{m}$) | 100 | 100 | 100 |
| Camera | K2 Summit | K2 Summit | K2 Summit |
| Detection mode | Super resolution | Super resolution | Super resolution |
| Pixel Size ($\text{\AA}/\text{Pixel}$) | 0.838 | 1.20 | 1.234 |
| Exposure (Sec) | 6.2 | 8.0 | 8.0 |
| Frame rate (Frames/sec) | 0.2 | 0.2 | 0.2 |
| Dose rate ($\text{e}^-/\text{\AA}^2/\text{frame}$) | 2.0 | 1.325 | 1.325 |
| Total dose ($\text{e}^-/\text{\AA}^2$) | 62.0 | 53 | 53 |
| Defocus range ($\mu\text{m}$) | 0.31 – 4.80 | 0.44 – 4.43 | 0.44 – 4.51 |
| Mean defocus ($\mu\text{m}$) | 1.29 | 1.50 | 2.24 |
| Reconstruction |  |  |  |
| Dataset | ssDNA | dsDNA | ssRNA |
| Micrographs | 2971 | 318 | 1591 |
| Particles extracted | 224,252 | 23,685 | 237,438 |
| Particles in final map | 101,990 | 21,864 | 23,924 |
| Box size ($\text{\AA}^3$) | 400<sup>3</sup> | 360<sup>3</sup> | 340<sup>3</sup> |
| Resolution ($\text{\AA}$) | 2.9 | 4.3 | 4.7 |
| Helical twist ($^\circ$) | 21.16 | 19.93 | 21.18 |
| Helical rise ($\text{\AA}$) | 3.17 | 2.92 | 3.17 |
| Model Validation |  |  |  |
| Dataset | ssDNA<br>PDB: 6E8G | dsDNA | ssRNA |
| EMRinger score | 2.92 | N/A | N/A |
| MolProbity score | 1.04 | N/A | N/A |
| Ramachandran favored (%) | 99.10 | N/A | N/A |
| Ramachandran allowed (%) | 0.90 | N/A | N/A |
| Ramachandran outliers (%) | 0.00 | N/A | N/A |
| Rotamer outliers (%) | 0.34 | N/A | N/A |
| C-beta deviations | 0.00 | N/A | N/A |
| Clashscore | 2.54 | N/A | N/A |
| RMS bonds ($\text{\AA}$) | 0.0069 | N/A | N/A |
| RMS angles ($^\circ$) | 1.16 | N/A | N/A |
| Mean B-factor | 52.5 | N/A | N/A |
| Maximum B-factor | 85.3 | N/A | N/A |
| Minimum B-factor | 31.6 | N/A | N/A |

As expected, density corresponding to the nucleic acid wraps around the interior of the cylindrical structure and tracks along the ‘groove’ between adjacent turns of the CHMP1B strand (**Figure 2A-B**). Although high-resolution features, including side chain densities, were visible for the protein subunits, the nucleic acid density was ‘smeared’ out into a featureless ribbon. This observation implies that the ssDNA does not follow the helical symmetry of the protein coat, as might be expected given the mixture of bases and the expected base repeat distance of ~3.1 Å for ssDNA (Mandelkern et al., 1981). Thus, the nucleic acid appears to interact via non-specific electrostatic interactions within the positively-charged surface of the copolymer lumen, making contacts that are not the same for each protein subunit (or DNA nucleotide). Much like the fluid and non-specific way in which membrane lipids would engage the structure, it appears that there is no specific register for the nucleic-acid oligomer along the IST1-CHMP1B copolymer surface, rendering ordered repeats undetectable. We also reconstructed dsDNA- and ssRNA-bound copolymer samples. These structures again showed well-ordered protein coats with smooth, featureless internal densities for the nucleic acids lining the groove between turns of the CHMP1B strand (**Table 1 and Figure 2-figure supplement 3**).

### Intersubunit connectivity between CHMP1B and IST1

The well-ordered ssDNA-bound IST1-CHMP1B filament sample allowed us to visualize new molecular details of the extensive inter-and intra-subunit interactions that were not evident in our previous nucleotide-free sample reconstructed at lower resolution (4 Å), (McCullough et al., 2015). The higher overal ionic strength (125 mM versus 25 mM) may have also increased the occupancy and order of C-terminal CHMP1B interacctions with IST1 (see below). The new maps reveal how CHMP1B makes specific contacts with *eight* neighboring monomers of CHMP1B in its fully extended conformation (**Figure 3A**). To facilitate descriptions of these interactions, we define a central CHMP1B molecule “i” (CHMP1B_i_) and its closest IST1 molecule “j” (IST1_j_), and then use integer steps to index neighboring molecules in the plus and minus directions (**Figure 3A-C** and **Figure 2-figure supplement 1C-D**). The largest intramolecular interface connects the CHMP1B_i_, and CHMP1Bi+1 molecules (and therefore also the CHMP1B_i-1_, and CHMP1B_i_ subunits) through hydrophobic contacts on α2, α3 and α4 on the CHMP1 B_i+1_ subunit with α1 and α4 on the CHMP1B_i_ subunit (**Figure 3D**). A second notable intrastrand contact is made with the CHMP1B_i+4_ subunit (and therefore also with the CHMP1B_i-4_ subunit), where the hydrophobic surface of CHMP1B_i_ α5 grasps the closed end of the α1/α2 hairpin from the CHMP1B_i+4_ subunit. This interaction appears to be a domain-swapped version of the equivalent *intramolecular* interaction seen beween helix α5 and the α2/α3 hairpin in the closed ESCRT-III conformation. IST1 intra-subunit connectivities are far less intricate, relying primarily on nearest-neighbor interactions along the helical path (i.e., IST_j_ interacts primarily with IST1_j+1_ and IST1_j-1_, **Figure 3E**). Residue I54 of the IST1_j_ subunit, near the closed end of the hairpin on helix α2, makes a key hydrophobic intersubunit contact with the IST1_j-1_ molecule. In the other direction, the IST1_j_ molecule presents D77 and R82 for favorable electrostatic inonic interactions with R55 and E57 of IST1_j+1_, respectively (**Figure 3E**). Finally, most of the intersubunit contacts between the two protein strands are salt bridges between residues from the C-terminal half of CHMP1B_i_, with complementary residues along the α1 helices of IST1_j_, IST1_j-1_, and IST1_j-2_ (**Figure 3F-G**).

**Figure 3:**
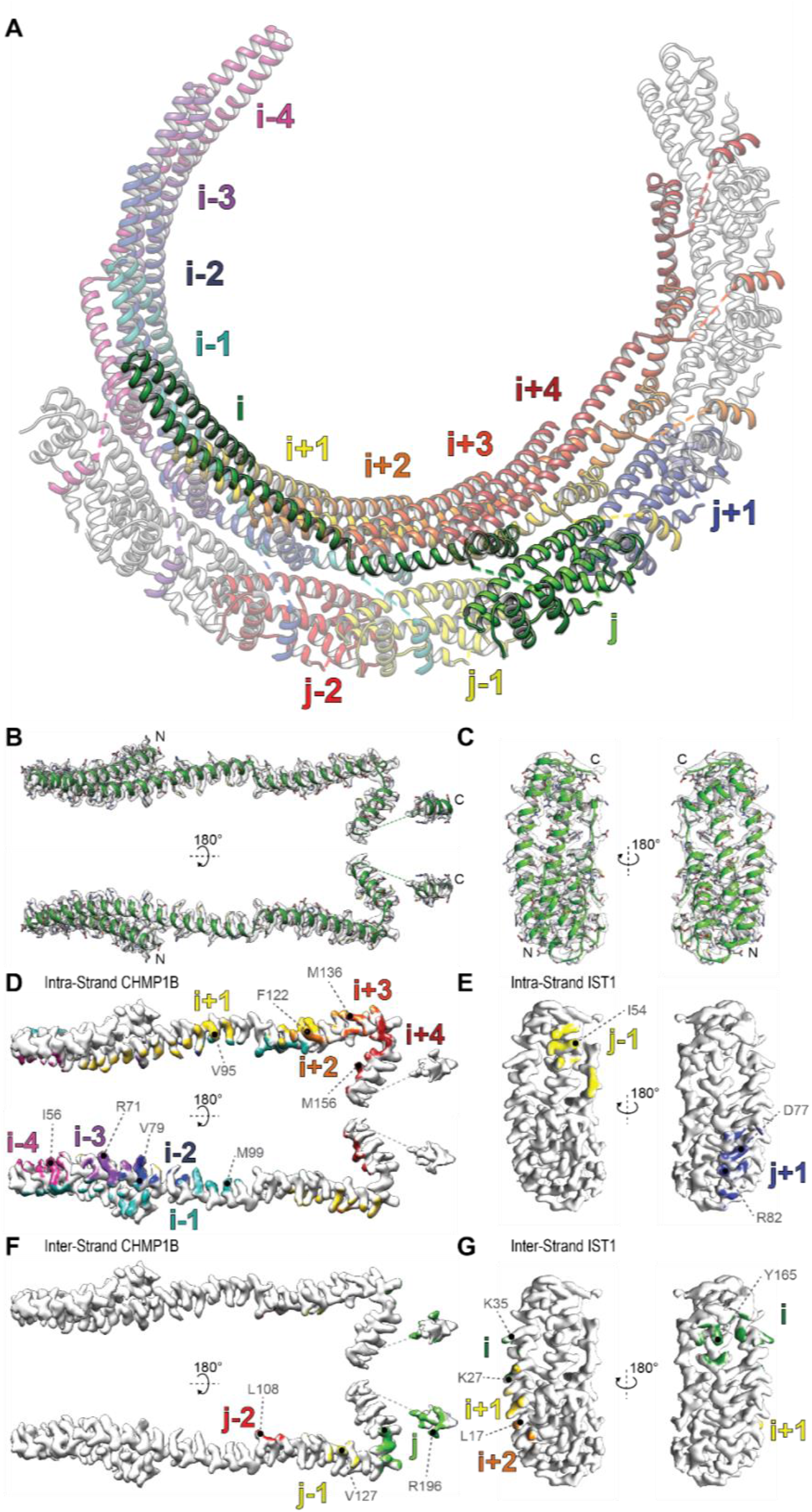
Intra- and inter-strand interactions within helical IST1-CHMP1B filaments. (**A**) Modeled ribbon diagram of the domain-swapped CHMP1B_i_ and IST1_j_ subunits within the assembled filament. The image shows a partial helical turn containing all of the subunits that contact the CHMP1B_i_ and IST1_j_ monomers. Modeled (**B**) CHMP1B and (**C**) IST1 fit within the segmented density corresponding to individual subunits within the assembled helical filaments. Note that all side chains are modeled. (**D**) and (**E**) Intra-strand interactions mapped onto surface renderings of the CHMP1B_i_ and IST1j subunits. (**F**) and (**G**) Inter-strand interactions mapped onto surface renderings of the CHMP1 Bi and IST1j subunits. Surface colors correspond to models in (**A**) and residues that make key contacts are labeled on their respective structures.

### CHMP1B MIM1 binds the outer surface of IST1

An important intermolecular contact that was not previously visualized is made between the MIM1-containing helix α6 of CHMP1B (residues 187–199) and the external surface of the IST1 subunit (**Figure 4A**). The MIM1 helix binding site is created by IST1 helices α1- α2, which buttress the MIM1 helix from below, and α5, which packs along one side of the CHMP1B MIM1 helix in an antiparallel orientation (**Figure 4B**). This interaction site may be partially occupied in the previous lower resolution maps of our protein-only IST1-CHMP1B filaments (McCullough et al., 2015) but the density is quite weak, perhaps owing to the low ionic strength conditions used in the previous reconstruction. Interestingly, an equivalent interaction was also reported previously in the crystal structure of yeast Ist1p bound to the isolated MIM1 helix from Did2p (the yeast ortholog of CHMP1B) (Xiao et al., 2009). The two structures overlay very well (Cα RMSD of <0.8 Å, **Figure 4C**), and alignment of the human C-terminal CHMP1B (187–199) and yeast Did2p (188–204) MIM1 helices shows that the contact residues are well conserved (**Figure 4A-C**). Our ability to position the C-terminal MIM1 of CHMP1B implies that a polypeptide chain spanning CHMP1B residues 165–186 must pass through the IST1 layer as it connects CHMP1B_i_ helix α5 to CHMP1B_i_ helix α6 (the CHMP1B MIM1 helix). The linker itself is not well defined in our structure, though the molecular connectivity is unambiguous owing to the short distance from the end of CHMP1B_i_ helix α5 to CHMP1B_i_ helix α6.

To investigate whether the CHMP1B MIM1 also binds IST1 in solution, we generated a series of N- and C-terminal truncations of CHMP1B and quantified their interactions with GST-tagged IST1_NTD_ using biosensor binding experiments. IST1_NTD_ adopts the closed conformation in solution (Bajorek et al., 2009b; Xiao et al., 2009) and remains closed in the assembled polymer (McCullough et al., 2015). The MIM1 binding pocket is therefore expected to be present in both structures. As shown in (**Figure 4-figure supplement 1A**), CHMP1B_4–199_(WT) bound weakly, but detectably to IST1_NTD_ (*K*_D_=92 μM), whereas a series of truncated CHMP1B proteins that lacked the MIM1 helix α6 did not exhibit detectable IST1_NTD_ binding (**Figure 4-figure supplement 1A**). Importantly, a peptide that corresponded to the CHMP1B MIM1 element alone (residues 186–199, denoted CHMP1Bα6) bound with full affinity (*K*_D_ = 91 μM). Thus, CHMP1B MIM1-IST1_NTD_ binding can be detected in solution, and this interaction accounts for all of the interaction energy between CHMP1B and IST1_NTD_ under conditions where the two proteins cannot copolymerize.

**Figure 4:**
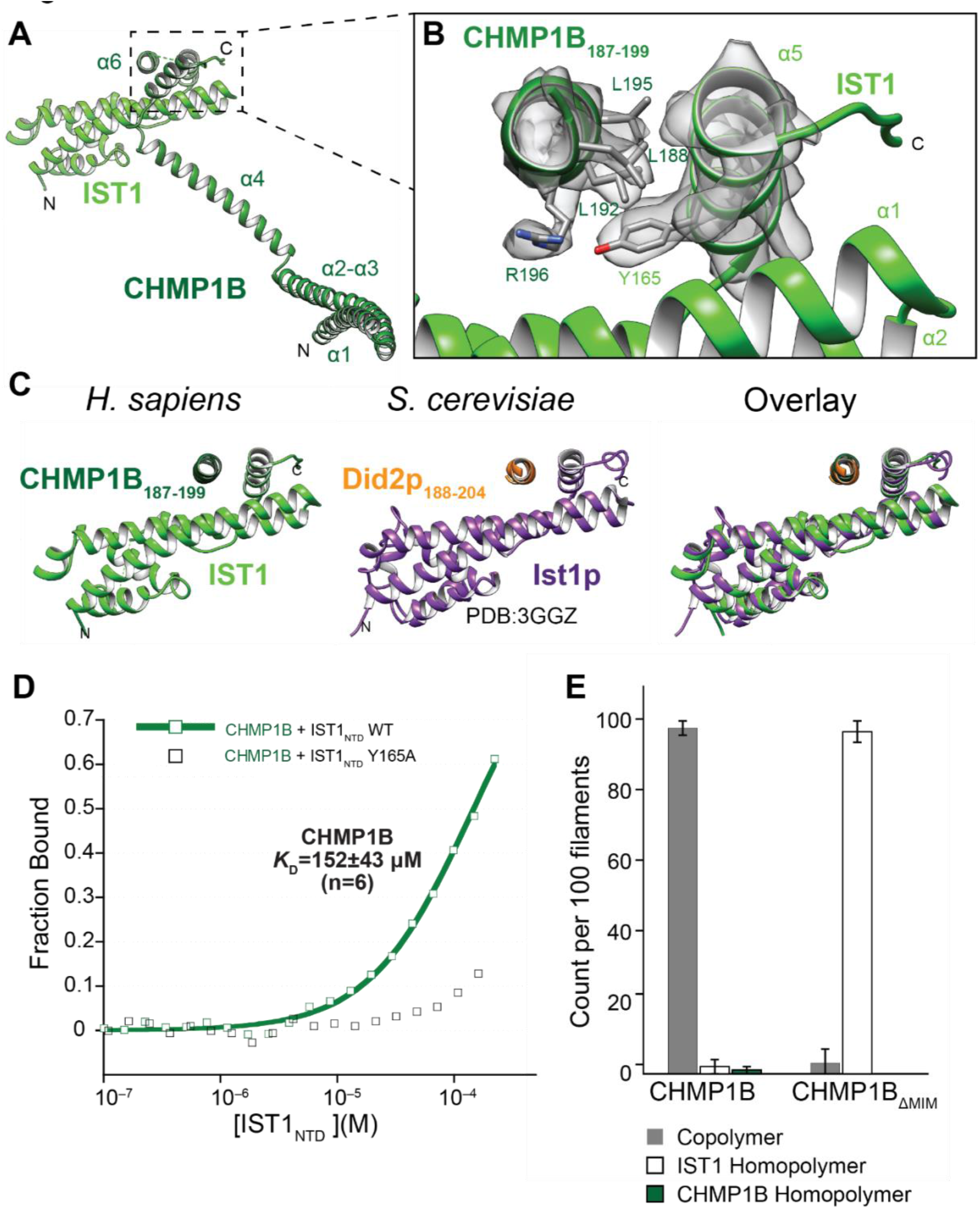
The CHMP1B MIM1 helix contacts the outer surface of IST1 and stabilizes the copolymer. (**A**) Ribbon diagram showing how the CHMP1B_i_ and IST1_j_ subunits interact. (**B**) Expanded view showing reconstructed density and fitted CHMP1B MIM1 and IST1 models. Note that the CHMP1 B MIM1 helix contacts a hydrophobic surface of IST1 created by the α1 - α2 hairpin and the perpendicular a5 helix. Key residues within the interface are labeled (Y165 for IST1 and L188, L192, L195 and R196 for CHMP1B). (**C**) Structural comparison of the human CHMP1B MIM1-IST1 interaction (Left) with the yeast Ist1p-Did2p MIM interaction (Middle) PDB:3GGZ (Xiao et al., 2009). (Right) Overlaid structures showing the high degree of similarity (Cα atom RMSD=0.76). (**D**) Fluorescence polarization binding isotherms for CHMP1B_169–199_ interacting with IST1_ntd_ WT (green squares) or IST1_ntd_ Y165A (black squares). Data points are averaged from at least three independent experiments. (**E**) Quantification of the filament type formed upon incubation of IST1 with either CHMP1B or CHMP1B _Δmim_ (CHMP1B_4–178_). Note that IST1 and CHMP1 B subunits can either copolymerize into double-stranded filaments (Copolymers) or homopolymerize (IST1 and CHMP1B Homopolymers, respectively)(McCullough et al., 2015). These different structures are readily distinguishable in negative stained EM images (see **Figure 4, supplemental figure 1**). In each case, 100 filaments were imaged and manually assigned to one of the three possible structures. Graphed values show the mean values from three replicate samples and error bars show standard deviations.

Mutational analyses were performed to test the specificity of the CHMP1B MIM1-IST1 interaction. Three leucine residues on one side of the amphipathic CHMP1B MIM1 helix α6 residues (L188, L192 and L195) appear to contact IST1 in our structure (**Figure 4B**). These residues were mutated to alanine in a pairwise fashion and IST1 binding was tested in the contexts of both full-length CHMP1B and the isolated CHMP1B MIM1 helix α6. In both contexts, the CHMP1B L188A/L192A and L192A/L195A mutations eliminated all detectable IST1 binding. (**Figure 4-figure supplement 1A**). Thus, CHMP1B and IST1 can interact in solution through the CHMP1B MIM1-IST1 interface seen within our copolymer structure. This interaction also nicely explains why CHMP1B binding was inhibited by a series of mutations along IST1 α2 (D64A, Y64D, and E67A) on the other side of the interface (Bajorek et al., 2009b).

### MIM1 binding to IST1 stabilizes the copolymer

To examine whether the CHMP1B MIM1-IST1 interaction also contributed to copolymer formation, we used the structure to identify another IST1 residue within the interface (Y165) and used fluorescence polarization anisotropy binding studies to demonstrate that the IST1 Y165A mutation eliminated CHMP1B MIM1 binding (**Figure 4D**). Importantly, the Y165A mutation also eliminated CHMP1B coassembly, as assessed by negative stain TEM (**Figure 4-figure supplement 1D**). Similarly, a CHMP1B construct that lacked the α5-α6 linker and the entire MIM1 helix α6 (CHMP1B_αi-α5_ or CHMP1B_ΔMIM_) (**Figure 4E** and **Figure 4-figure supplement 1C**) was also severely compromised (although not entirely lacking) in the ability to copolymerize with IST1 (**Figure 4E** and **Figure 4-figure supplement 1B and 1D**). Hence, CHMP1B MIM1 helix binding to the outer surface of IST1 significantly stabilizes the helical double stranded filament.

We next tested whether IST1 can bind the C-terminal tails of other ESCRT-III proteins, especially those with related MIM1 sequences. A collection of C-terminal labelled peptides from all human ESCRT-III proteins was generated (**Table 2**) and tested for binding to IST1_NTD_ using fluorescence polarization anisotropy (**Figure 5A**). In this initial screen, IST1_NTD_ binding was only detectable for the MIM1 helix of CHMP1B. This observation was surprising because the CHMP1B MIM1 helix is highly similar in sequence to its closest ESCRT-III homolog, CHMP1A (**Figure 5B and Figure 5-figure supplement 1A**). We therefore tested IST1_NTD_ binding to a shorter CHMP1A MIM1 construct that matched exactly the length of the CHMP1B MIM1 peptide, as well as a control CHMP2A MIM1 peptide of the same length. In this context, the CHMP1A and CHMP1B peptides bound IST1_NTD_ with similar affinities, whereas the CHMP2B peptide did not bind detectably (**Table 2, Figure 5C**). These observations indicate that IST1 binds weakly but specifically to the C-terminal MIM1 tails of both CHMP1A and CHMP1B, and suggests that upstream CHMP1A sequence elements may autoinhibit or occlude this interaction.

**Figure 5:**
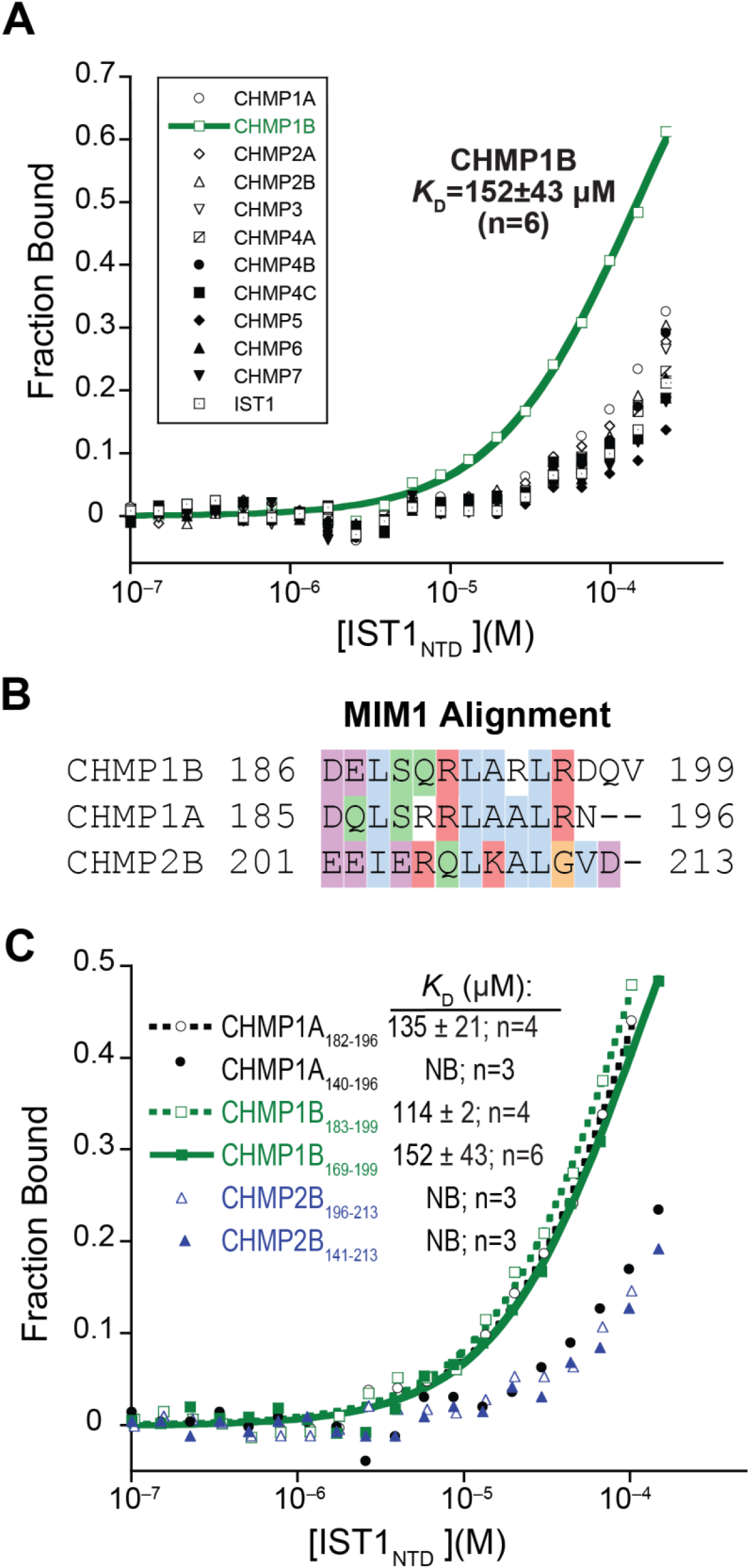
IST1_ntd_ binds specifically to the MIM1 helices of CHMP1A and CHMP1B. (**A**) Fluorescence polarization binding isotherms for IST1_ntd_ and the indicated ESCRT-III C-terminal peptides. Curves shown represent the averages of at least three independent experiments. Binding isotherms for all ESCRT-III proteins except CHMP1B_169–2199_ ***(Kd*** = 152 ± 43 μM) correspond to dissociation constants greater than 400 μM. (**B**) Protein sequence alignment highlighting chemically similar residues for CHMP1B, CHMP1A, and CHMP2B MIM1 helices. (**C**) Fluorescence polarization binding isotherms for IST1ntd interactions with the indicated ESCRT-III C-terminal peptides. The experiment shows that IST1_ntd_ binds specifically to the CHMP1A and CHMP1 B MIM1 helices, and that a minimal CHMP1A MIM1 helix binds IST1_ntd_ better than a longer CHMP1A construct. Binding curves correspond to averages from the numbers of experiments indicated in the inset legend.

**Table 2:**
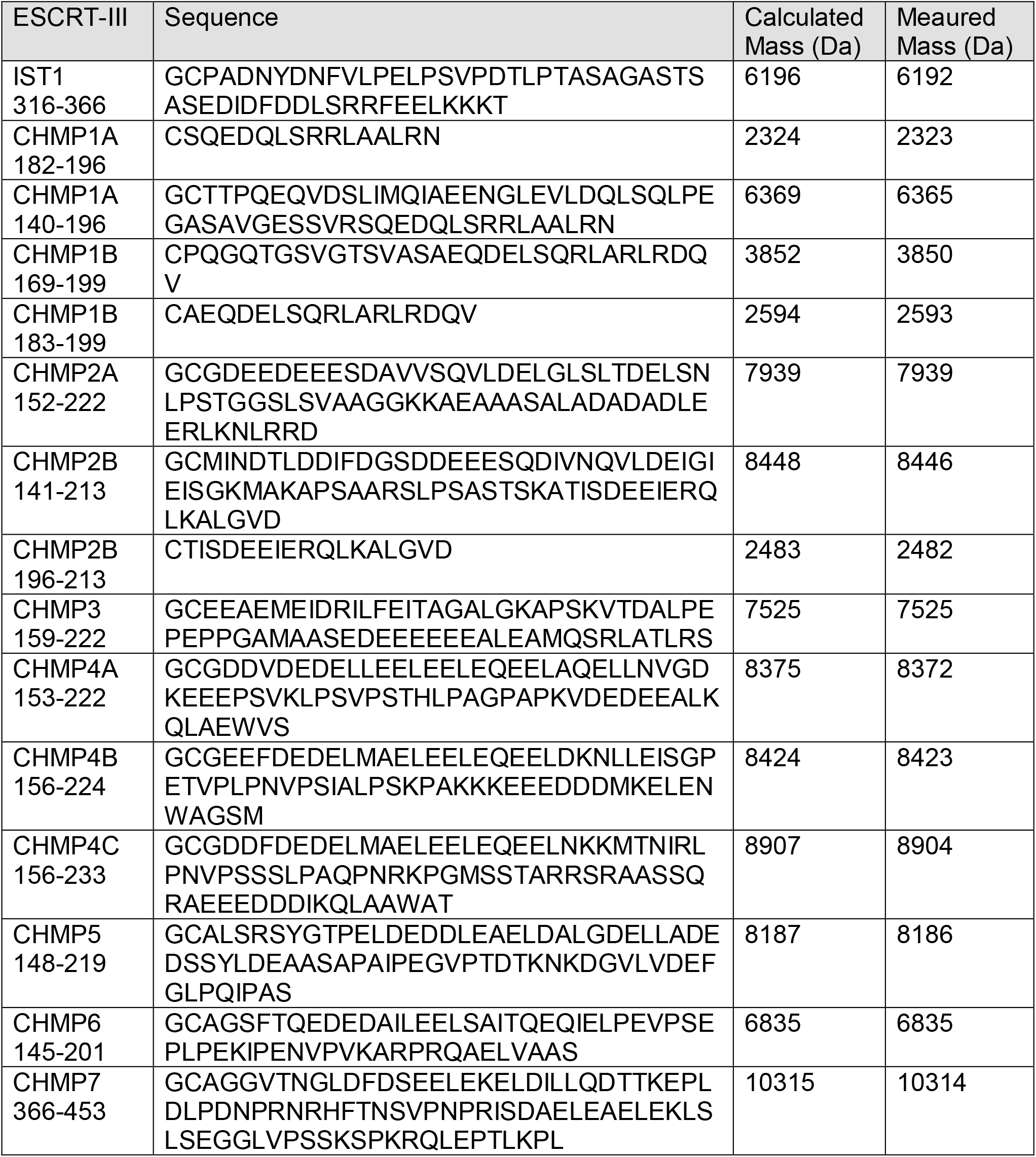
ESCRT-III C-terminal fluorescent polarization anisotropy petpides.

**Table 3:** Plasmid deposition information.

| Plasmid | Internal ID | DNASU # or Addgene # | UniProt # |
| --- | --- | --- | --- |
| pET16b IST1 1-366 | WISP 07-72 | HsCD00671538 | P53390-4 |
| pGEX IST1 1-189 | WISP 07-74 | HsCD00520979 | P53390-4 |
| pGEX IST1 1-189 Y165A | WISP 09-55 | HsCD00520956 | P53390-4 |
| pGEX CHMP1B 4-199 | WISP 08-221 | HsCD00520966 | Q7LBR1 |
| pCA528 CHMP1B 4-199 | WISP 18-23 | 115325 (Addgene) | Q7LBR1 |
| pCA528 CHMP1B 1-199 | WISP 18-24 | 115326 (Addgene) | Q7LBR1 |
| pGEX CHMP1B 4-178, aka CHMP1B<sub>ΔMIM</sub> | WISP 09-279 | 108284 (Addgene) | Q7LBR1 |
| pGEX CHMP1B 186-199 | WISP 09-283 | 108285 (Addgene) | Q7LBR1 |
| pGEX CHMP1B 4-143 | WISP 09-278 | 108286 (Addgene) | Q7LBR1 |
| pGEX CHMP1B 144-178 | WISP 09-281 | 108287 (Addgene) | Q7LBR1 |
| CHMP1B 186-199 L188A/L192A | WISP 09-286 | 108288 (Addgene) | Q7LBR1 |
| CHMP1B 186-199 L192A/L195A | WISP 09-287 | 108290 (Addgene) | Q7LBR1 |
| CHMP1B 4-199 L188A/L192A | WISP 09-284 | 108289 (Addgene) | Q7LBR1 |
| CHMP1B 4-199 L192/L195A | WISP 09-285 | 108291 (Addgene) | Q7LBR1 |
| pCA528 CHMP1A 140-196 | WISP18-1 | 104593 (Addgene) | Q9HD42 |
| pCA528 CHMP2A 152-222 | WISP18-3 | 104594 (Addgene) | O43633 |
| pCA528 CHMP2B 141-213 | WISP18-4 | 104595 (Addgene) | Q9UQN3 |
| pCA538 CHMP3 159-222 | WISP18-5 | 104599 (Addgene) | Q9Y3E7 |
| pCA528 CHMP4A 153-222 | WISP18-6 | 108256 (Addgene) | Q9BY43 |
| pCA528 CHMP4B 156-224 | WISP18-7 | 104596 (Addgene) | Q9H444 |
| pCA528 CHMP4C 156-233 | WISP18-9 | 108257 (Addgene) | Q96CF2 |
| pCA528 CHMP5 148-219 | WISP18-10 | 108259 (Addgene) | Q9NZZ3 |
| pCA528 CHMP6 145-201 | WISP18-11 | 104597 (Addgene) | Q96FZ7 |
| pCA528 CHMP7 366-453 | WISP18-12 | 104598 (Addgene) | Q8WVX9 |
| pCA528 IST1 316-366 | WISP14-121 | HSCD00696258 | P53390-4 |

## DISCUSSION

Our studies show that the IST1-CHMP1B copolymer can encapsulate and protect nucleic acid oligomers, and that copolymer formation is nucleated by ssRNA, ssDNA and dsDNA (**Figure 1**). In each case, the nucleic acid polymer winds along the strongly basic ‘groove’ formed at the interface between adjacent turns of the inner-strand protein, CHMP1B (**Figures 2 and 3**). Future work will assess whether CHMP1B and IST1 (or CHMP1B alone) bind nucleic acids in living cells. As noted earlier, published observations suggest that ESCRT-III proteins, including CHMP1A, have poorly understood roles that appear to involve nuclear localization and/or extra-nuclear co-localization with nucleic acids. Any such role would necessarily be independent of membrane-remodelling roles, however, because the same lumenal surface of CHMP1B—defined primarily by the highly charged surface of helix αl that faces the center of the helical tube—is used for both membrane and nucleic-acid binding.

Interestingly, human CHMP1B is an intron-less retrogene that arose in a common ancestor of the placental mammals via copying and reinsertion of a primordial CHMP1B gene (Nels Elde and Diane Downhour, personal communication and (Ciomborowska et al., 2013)). The fate of the original CHMP1B gene in placental mammals varies by species, but in all primates the parental copy is now a pseudogene. Expansion of the CHMP1 gene family and retro copying of CHMP1B in placental mammals may have allowed CHMP1B to evolve new functions, either as an inverted outside-in-oriented ESCRT-III, or more speculatively to sense and coat nucleic acids. By contrast, the gene encoding mammalian CHMP1A is not a retrogene and has also been reported to play a role in the biogenesis of a specialized MVB that gives rise to secretory extracellular vesicles (Coulter et al., 2018).

Our cryoEM structure of a full-length IST1-CHMP1B complex with ssDNA is also of value in providing the most detailed view to date of an ESCRT-III filament assembled from essentially full-length proteins. The 2.9 Å resolution reconstruction revealed that the C-terminal tails from the inner CHMP1B strands unexpectedly extend to the filament exterior, where their MIM1 helices bind the exterior surface of IST1 and interlock the two strands even more extensively than originally appreciated (**Figure 3**). This interaction is an essential component of the assembled filament, because its removal impairs filament formation (**Figure 4**). The specificity of this interaction (**Figure 5**), and the conservation of analogous interactions between yeast Ist1p with Did2p and human IST1 with both human CHMP1 proteins (**Figure 4**) helps explain why IST1 and CHMP1A/B proteins commonly function together as an ESCRT sub-module (Xiao et al., 2009).

Exposure of the C-terminal CHMP1B MIM1 element on the tube exterior also has important implications for understanding how ESCRT-III filaments recruit MIT domain-containing proteins that function as cofactors, particularly the AAA ATPases that disassemble ESCRT-III filaments and other biopolymers. Previous work has shown that the CHMP1B MIM1 element binds the MIT domain of SPASTIN, and that disruption of this interaction impairs MVB sorting and cytokinetic abscission (Xiao et al., 2009; Yang et al., 2008). Our structure now reveals that the CHMP1B MIM1 element is available to help link IST1, CHMP1B and SPASTIN into a physiologically relevant protein interaction network (Allison et al., 2017; Allison et al., 2013; Bajorek et al., 2009a; McCullough et al., 2015; Reid et al., 2005; Xiao et al., 2009; Yang et al., 2008). Indeed, the C-terminal MIM1 helix of CHMP1B can bind the MIT domains from a series of different proteins, including VPS4 (Stuchell-Brereton et al., 2007), LIP5, a regulator of VPS4 activity (Shim et al., 2008), UBPY, a ubiquitin isopeptidase (Row et al., 2007) and SPASTIN (Yang et al., 2008). Comparison of our structure with the crystal structure of the SPASTIN MIT-CHMP1B MIM1 complex (Yang et al., 2008) reveals that the same CHMP1B MIM1 surface is used to bind both IST1 and the SPASTIN MIT domain. Hence, the CHMP1B MIM1 element could not bind both proteins simultaneously, and would instead need to release from the IST1 surface in order to engage SPASTIN (Figure 6). This situation creates multiple opportunities for conformational evolution and regulation because the IST1-CHMP1B filaments will be most stable when the CHMP1B MIM1 element is bound to the IST1 surface. Releasing this interaction will cost energy and competitively attenuate binding by the VPS4 and SPASTIN MIT AAA ATPases. However, when MIT domains do bind the CHMP1B MIM element, these interactions will tend to destabilize and perhaps prime IST1-CHMP1B filaments for remodeling or disassembly.

**Figure 6:**
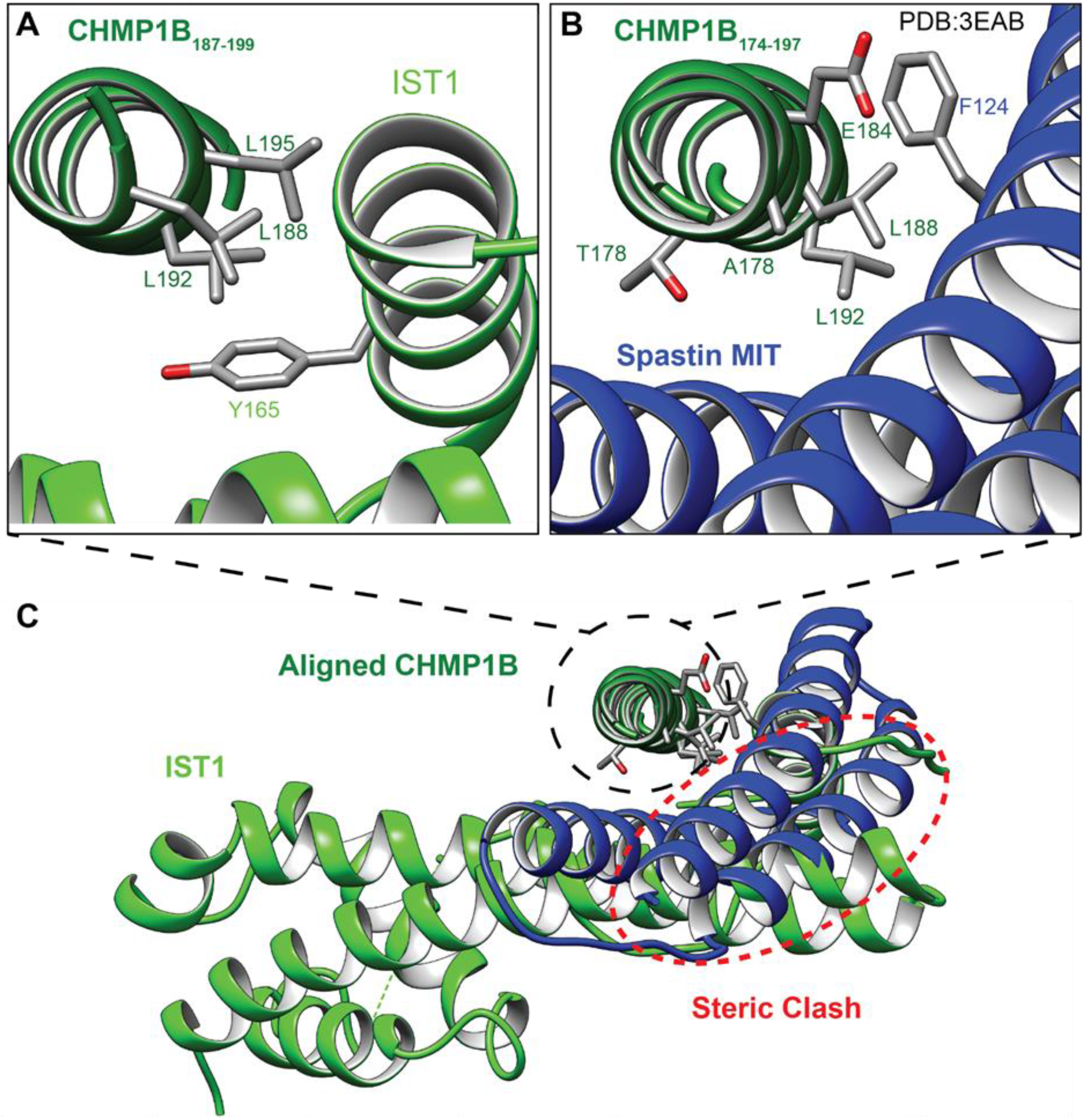
Spastin MIT domain binding is mutually exclusive with IST1 for CHMP1B MIM1 helix binding. (**A**) Structure of the CHMP1B MIM1 helix bound to IST1 within IST1-CHMP1B copolymeric filaments. (**B**) Crystal structure of the extended CHMP1B MIM1 helix bound to the SPASTIN MIT domain (PDB: 3EAB) (Yang et al., 2008). Residues shown to be important for the CHMP1B MIM1-IST1 interaction (this study) or for the CHMP1B MIM-SPASTIN MIT interaction (Yang et al., 2008) are shown in stick representations. (**C**) Superposition of the CHMP1 B MIM1 helix (dark green) in complex with IST1 (light green) or SPASTIN MIT (blue, PDB: 3EAB) (Yang et al., 2008). The two complex structures were aligned on their CHMP1 B MIM 1 helices (black dashed circle). A region of major steric clash (red dashed oval) indicates that the CHMP1 B MIM1 helix cannot bind simultaneously to IST1 and SPASTIN.

Finally, our structure raises the possibilty that other pairs of ESCRT-III proteins may also interact via their MIM elements. The inter-ESCRT-III MIM1 binding pocket of IST1 is specific for Did2p/CHMP1A/B-like helical tails (**Figure 5**), but we speculate that ESCRT-III proteins, like CHMP2, CHMP3, and CHMP4 subunits, could recognize and co-polymerize with their preferred binding partners via analogous MIM interactions between closed (donor) and open (acceptor) subunits. In this regard, it is important to note that many ESCRT-III structural studies published to date have employed N-terminal "core" constructs (Bajorek et al., 2009b; McCullough et al., 2015; McMillan et al., 2016; Muziol et al., 2006; Tang et al., 2015). These constructs have the advantage that removing C-terminal sequences typically removes autoinhibitory elements, and thereby favors conversion of the closed to open coformation and promotes polymerization. Our IST1-CHMP1B structure reveals, however, that C-terminal MIM helices can also contribute specific intermolecular interactions that stabilize this, and perhaps other, ESCRT-III polymers.

## Figures and Legends

**Figure 1 – figure supplement 1:**
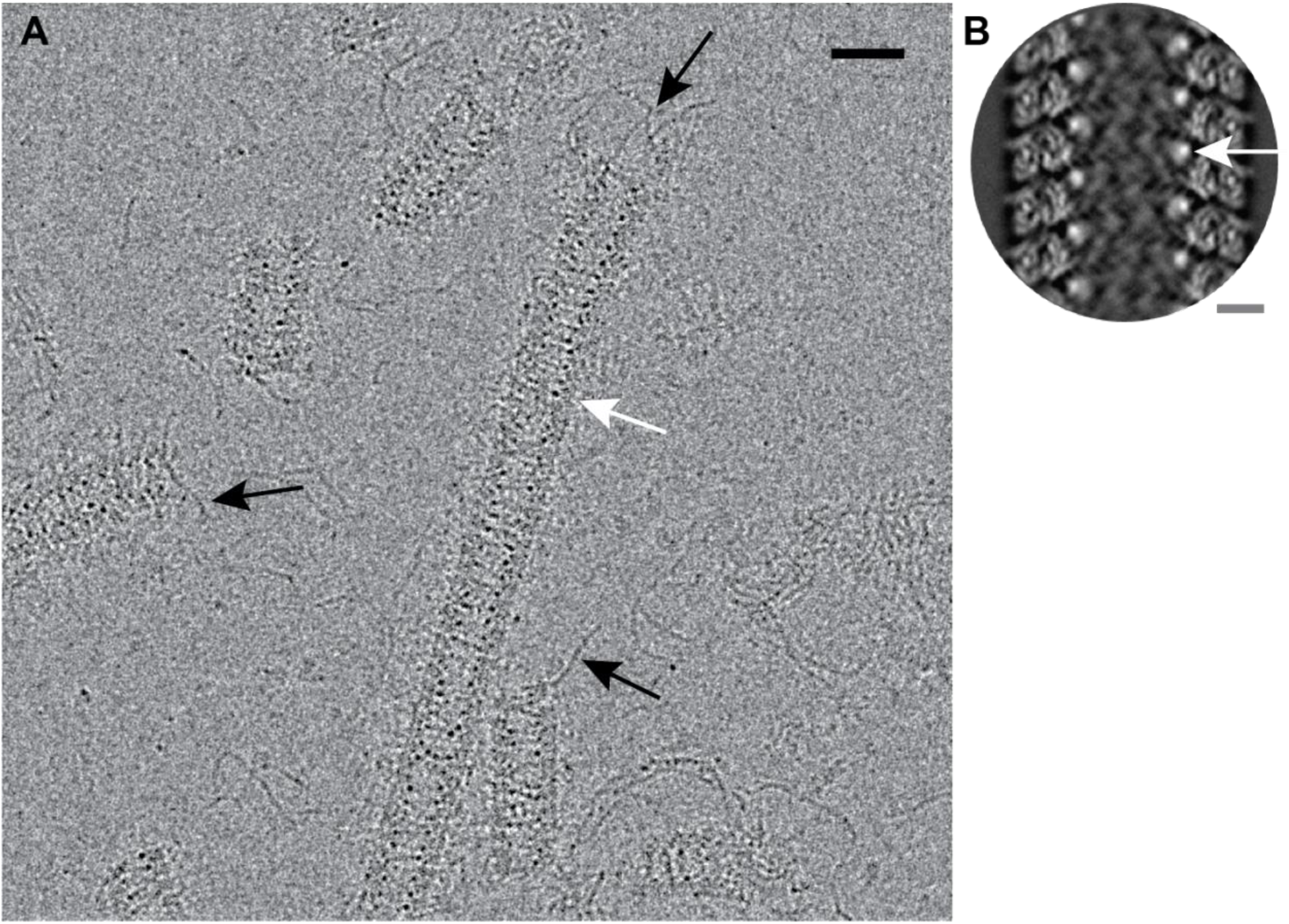
CryoEM image of ssDNA-templated IST1-CHMP1B copolymers. (**A**)Electron cryo-micrograph of the IST1-CHMP1B copolymer assembly bound to ssDNA. Note that ssDNA can be visualized as dark densities within the lumen of the protein filaments (white arrow) and extruding from the ends of the helical tubes of filaments (black arrows). Scale bar: 25 nm. (**B**) A 2D class average of the IST1-CHMP1B filament with visible density corresponding to ssDNA within the groove formed between adjacent turns of the helical density (white arrow). Scale bar: 5 nm.

**Figure 2 – figure supplement 1:**
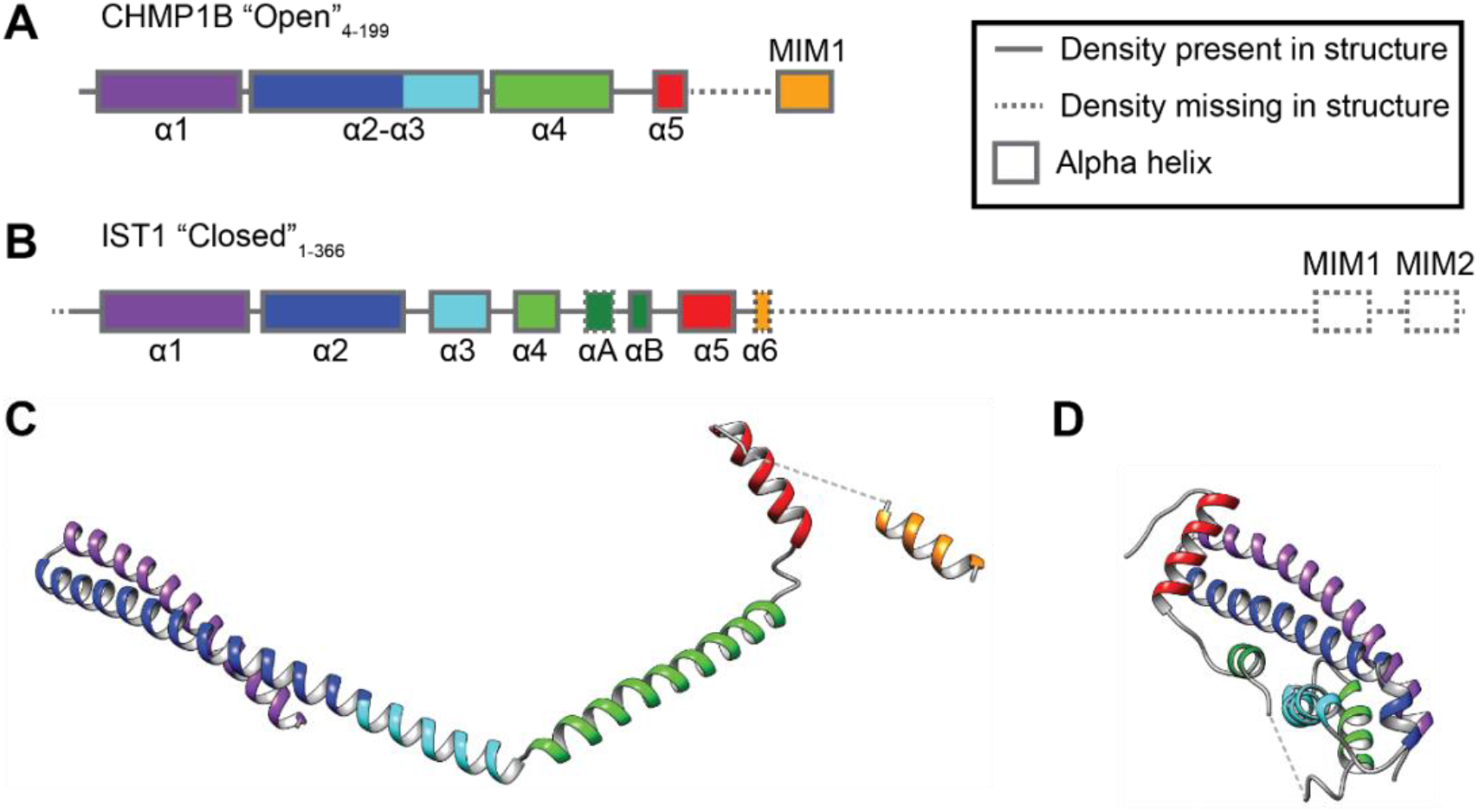
Secondary structures of the CHMP1B and IST1 monomers. Secondary structure diagram for: (**A**) CHMP1B and (**B**) IST1 proteins as modelled within the reconstruction. Dashed line segments indicate missing density in the reconstruction, despite the presence of full-length CHMP1B and IST1 in the reconstructed sample. (**C**) and (**D**) Models of CHMP1B and IST1 structures. The structures are aligned on the helix α1-α2 hairpin, and α-helical segments are colored to match the corresponding secondary structure diagrams in (A) and (B).

**Figure 2 – figure supplement 2:**
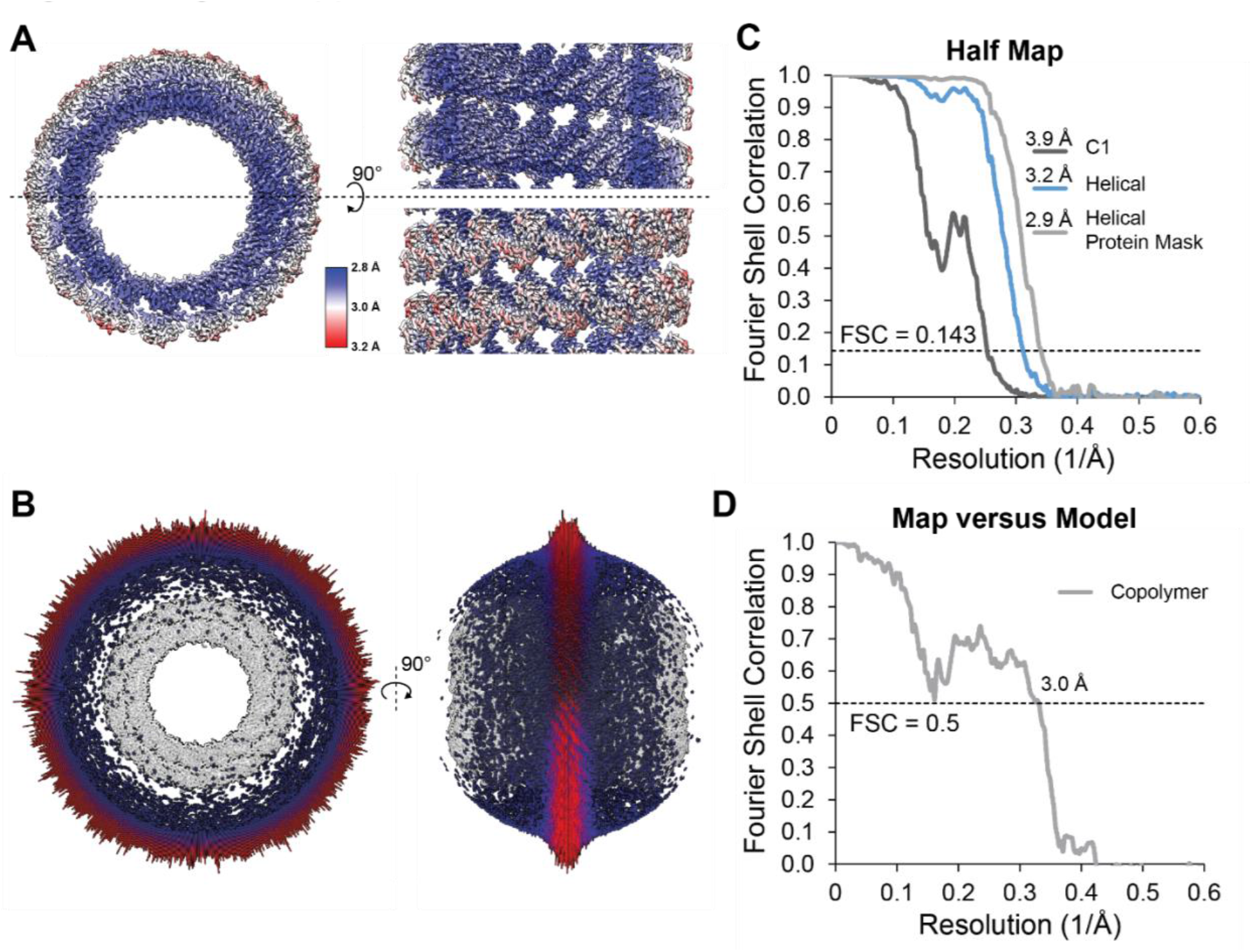
Resolution estimates and model validation for the IST1-CHMP1B-ssDNA reconstruction. (**A**) LocalRes rendered view of the local resolution of the CHMP1B and IST1 regions from the reconstructed IST1-CHMP1B-ssDNA complex, labelled from 3.2 Å (red) to 2.8 Å (blue). End on view (left), lumenal view (upper right) and exterior view (lower right). (**B**) Orthogonal views of the angular distributions of particles for the IST1-CHMP1 B-ssDNA reconstruction. (**C**) Half map FSC curves for the asymmetric spherical mask (C1 - dark grey), helical symmetry imposed with a featureless cylindrical mask (Helical - light blue), and helical symmetry imposed with a soft protein mask (Helical Protein Mask - grey) reconstructions. (**D**) Map versus model FSC for both layers of the copolymer (grey). The comparison was made using 36 subunits for each double-stranded protein layer (two full turns of the helix).

**Figure 2 – figure supplement 3:**
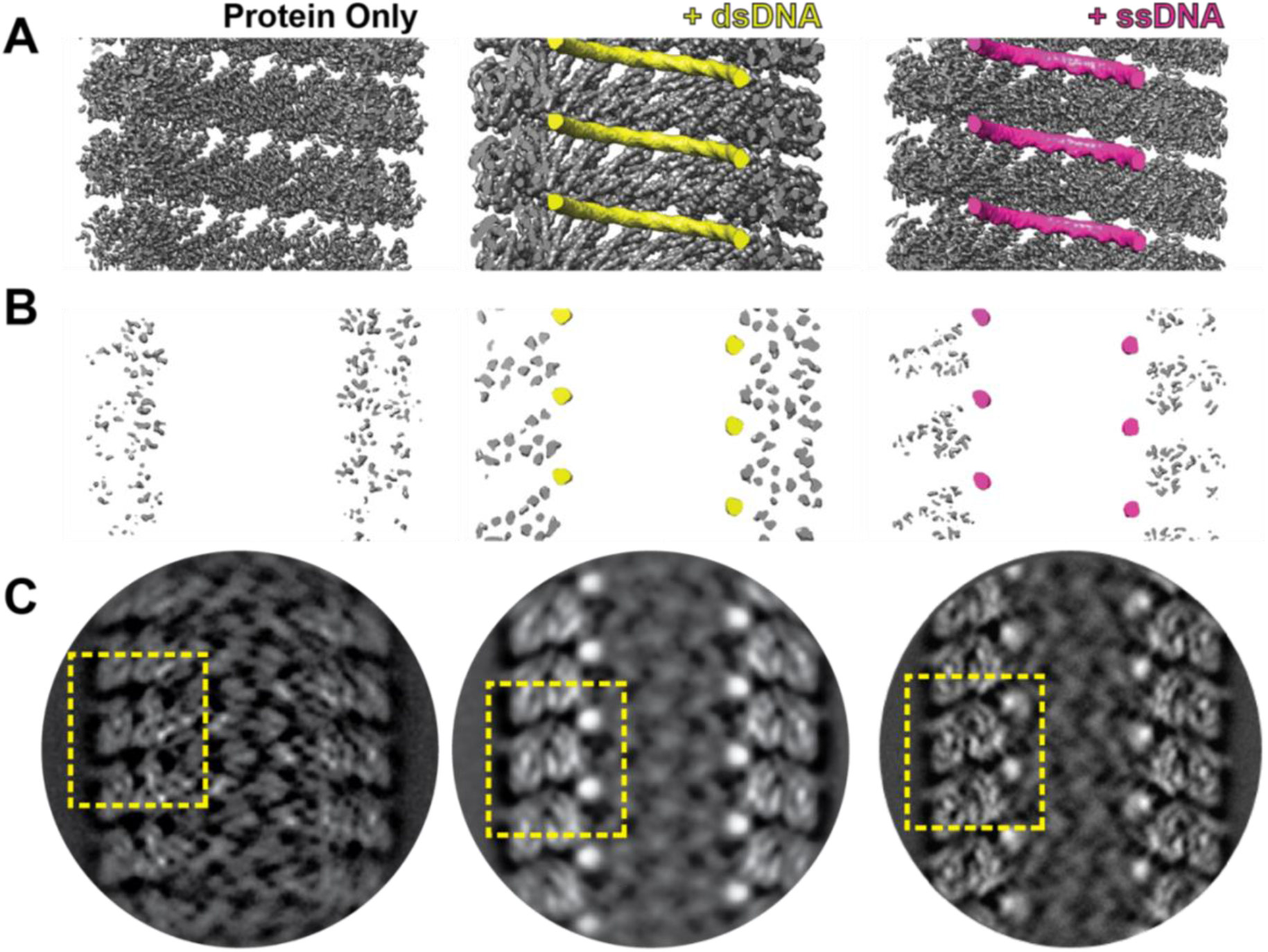
ssDNA and dsDNA occupy the same nucleic-acid binding groove between adjacent turns of the inner CHMP1B strands. (**A**) Internal cutaway views of different nucleotide-templated IST1-CHMP1B copolymers. Left to right: Protein only structure (determined previously, EMDB:6461 (McCullough et al., 2015)), and structures in complex with dsDNA and ssDNA. (**B**) Cross-sectional view highlighting the positions of the polynucleotides within the helical assemblies. (**C**) Reference-free 2D class averages from the corresponding copolymer datasets.

**Figure 4 – figure supplement 1:**
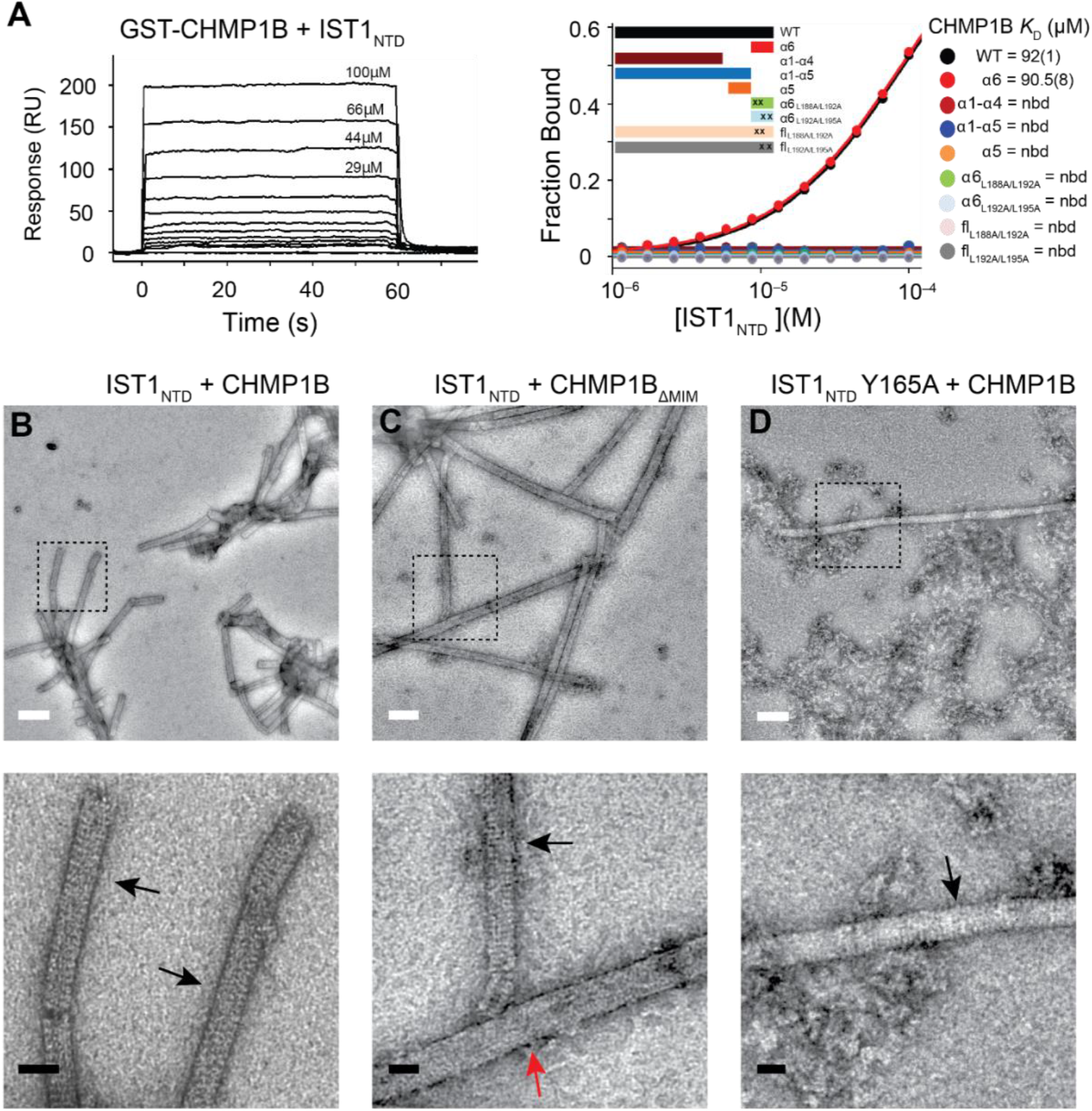
The CHMP1B MIM1 helix binds IST1 specifically and stabilizes the CHMP1B-IST1 copolymer. (**A**) (Left) Representative SPR sensorgram for the indicated concentrations of IST1_NTD_ binding to immobilized GST-CHMP1B. (Right) Biosensor binding isotherms for the indicated immobilized GST-CHMP1B constructs binding to IST1_NTD_. Dissociation constants are reported from one experiment, with fitting errors from a 1:1 binding model shown in parenthesis. (**B-D**) Reactions between the designated IST1_NTD_ and the designated CHMP1B proteins, imaged by negative stained EM at low magnification (upper panels, white scale bars are 100 nm) or medium magnification images of the boxed regions of the corresponding upper panel (lower panels, black scale bars are 25 nm). (**B**) Wild type IST1_NTD_ with wild type CHMP1B. Note that only CHMP1B-IST1_NTD_ copolymers were observed in this reaction (highlighted by black arrows in the lower panel). (**C**) Wild type IST1_NTD_ with a mutant CHMP1B construct lacking the MIM1 helix (CHMP1_ΔMIM_). Note that the vast majority of the filaments formed were IST1 homopolymers (red arrow) but CHMP1B-IST1_NTD_ copolymers (black arrow) were also occasionally observed. (**D**) IST1_NTD_ Y165A mutant with wild type CHMP1B. This reaction predominantly generated protein aggregates, but occasionally produced copolymeric IST1_NTD_ Y165A-CHMP1B filaments (black arrow).

**Figure 5 – figure supplement 1:**
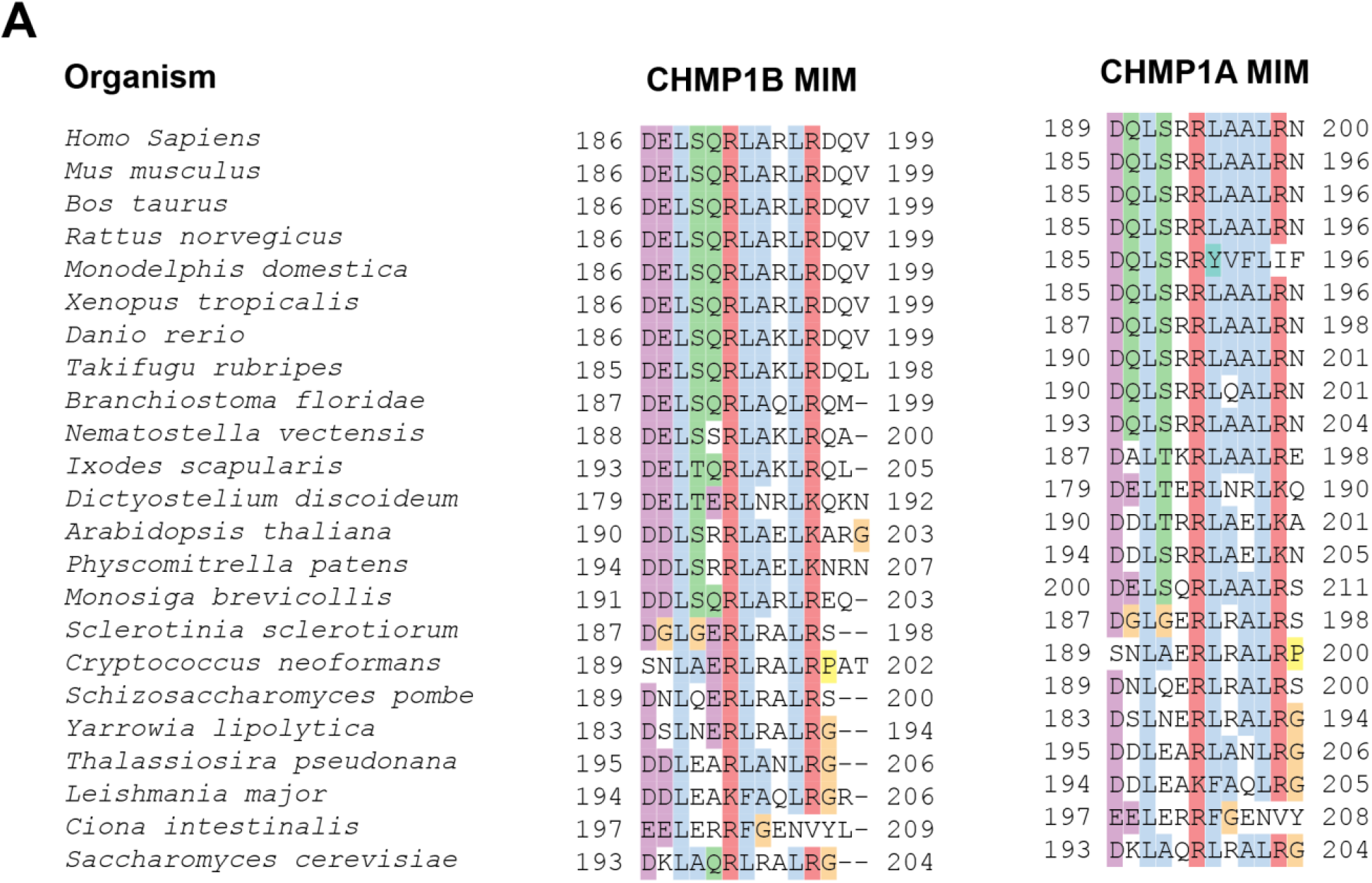
Conservation between CHMP1B and CHMP1A MIM1 Helices. (**A**) CHMP1B and CHMP1A MIM1 helix sequence alignments, highlighting the conservation of chemically similar residues.

## Materials and Methods

### Purification of IST1 and CHMP1B proteins

The purification of IST1 FL (McCullough et al., 2015), IST1_1–189_ (IST1_NTD_) and CHMP1B_4–199_ (Bajorek et al., 2009a) have been described previously. 1ST1_1–189_ Y165A was purified following the protocol for 1ST1_1–189_ (Bajorek et al., 2009a) and the yield was < 30 mg from a 6 L culture. IST1_1–189_ Y165A protein identity was confirmed by mass spectrometry (MrCalc= 21643.67 Da, Mr_Exp_= 21643.66 Da). CHMP1B_4–178_ was purified following the protocol for CHMP1B4–199 (Bajorek et al., 2009a) and the yield was < 2 mg from a 6 L culture. CHMP1B_4–178_ protein identity was confirmed by mass spectrometry (Mr_Calc_ = 19617.90 Da, MrExp= 19617.88 Da). CHMP1B_4–199_ for structure determination was cloned into a His-SUMO fusion to have an N-terminus with no residual amino acids in contrast to the pGEX expressed proteins that contain an extra Gly-His appended to the N-terminus after TEV cleavage. This His-SUMO-CHMP1B_4–199_ was expressed in BL21-Codon Plus (DE3) RIPL cells (Agilent, Santa Clara, CA, USA) in ZYP-5052 auto-induction media (Studier, 2005). Cells were harvested and frozen at -80 °C. All subsequent steps were performed at 4 °C unless otherwise noted. Thawed cells were resuspended in lysis buffer (50 mM Tris, pH 8 at 4 °C, 300 mM NaCl, 10 mM Imidazole, 1 mM DTT, 0.5 mM EDTA) and supplemented with lysozyme (0.2 mg/mL), protease inhibitors (PMSF, pepstatin, aprotinin, leupeptin), DNAse I (Roche, Germany), and 2 mM MgSO_4_. Each gram of cell paste was suspended in four mL of lysis buffer and cells ruptured using freeze-thaw followed with emulsification using Emulsiflex (Avestine, Inc., Canada). Lysate was further processed with addition of 0.125% sodium deoxycholate, centrifuged at 35,000 *g* for 1 h, and the supernatant filtered using a 0.45 μm membrane. Cleared lysate was loaded onto 10 mL of cOmplete His-Tag purification resin (Roche, Germany), incubated for 1 hour, and washed extensively with lysis buffer. The fusion protein was eluted from the resin with 300 mM imidazole in lysis buffer and placed in a 12.5 kDa dialysis membrane bag with His-tagged ULP1 protease (0.75 mg), for removal of the His-SUMO tag in dialysis buffer (50 mM Tris pH 8.0 at 4 °C, 150 mM NaCl, 1 mM DTT). Following quantitative cleavage of the fusion protein, the solution was applied to a fresh cOmplete resin to bind His-SUMO tag and His-tagged ULP1 and CHMP1B_4–199_ was collected in the flowthrough. CHMP1B_4–199_ was further purified by Superdex-75 size exclusion chromatography (GE Healthcare Life Sciences, USA). Typical protein yields were 5–10 mg/L of culture. His-SUMO-CHMP1B_1–199_ can also be purified using the same purification protocol as His-SUMO-CHMP1 B_4–199_ (data not shown).

### Purification of ESCRT-III C-terminal peptides

ESCRT-III C-terminal peptides were expressed as His-SUMO-fusion proteins (except for CHMP1 B_169–199_, CHMP1A_182–196_, CHMP1B_183–199_, and CHMP2B_196–213_ which were synthesized by the University of Utah DNA/Peptide Synthesis Core) and were each expressed in 2 L cultures of BL21-Codon Plus (DE3) RIPL cells (Agilent, Santa Clara, CA, USA) in ZYP-5052 auto-induction media (Studier, 2005). All purification steps were performed at 4°C. Cells expressing ESCRT-III C-terminal fragments were lysed by sonication in buffer (40 mL/L of culture) containing 50 mM Tris, pH 7.2, 150 mM NaCl, 5 mM imidazole, 2 mM DTT, 0.5 mM EDTA, and 0.125% sodium deoxycholate, supplemented with lysozyme, protease inhibitors, and DNAse I (Roche, Germany). Clarified cell lysate was incubated with 10 mL of cOmplete His-Tag purification resin (Roche, Germany) for 20 min, washed with 10 column volumes of wash buffer: 50 mM Tris, pH 7.2, 500 mM NaCl, 5 mM Imidazole, 5 mM DTT, 0.5 mM EDTA followed by 10 column volumes of wash buffer containing 150 mM NaCl. The MIM peptides were cleaved from the HIS-SUMO affinity tag and eluted from the resin by overnight inucbation and cleavage with ULP1 protease (0.7 mg) in 40 mL of the 150 mM NaCl wash buffer at 4°C. The cleaved ESCRT-III C-terminal peptides were collected from the column flow through and dialyzed against Q-sepharose-binding buffer (25 mM NaPi, pH 6.5, 50 mM NaCl, 2 mM DTT, 0.5 mM EDTA) for further purification by Q-sepharose anion exchange chromatography (GE Healthcare Life Sciences, USA) with a linear gradient from 50 mM to 1 M NaCl. Fractions containing peptides were pooled and dialyzed against gel filtration buffer (25 mM Tris, pH 7.2, 150 mM NaCl, 1 mM TCEP, 0.5 mM EDTA) and further purified by Superdex-75 size exclusion chromatography (GE Healthcare Life Sciences, USA). Typical peptide yields were 4.5 mg/L culture (IST1). Purified ESCRT-III C-terminal fragments contain non-native ‘GC’ residues at their N-termini, respectively, and masses were confirmed either before labeling (CHMP4C, CHMP1A_140–196_, CHMP1B_143–199_) or after labeling (all other peptides; +Dye yields a mass shift of 463.4 Da) by mass spectrometry (see **Table 2**).

### Fluorescent labeling of peptides

Fluorescent labeling was performed by the University of Utah DNA/Peptide Synthesis Core as described previously (Caballe et al., 2015). Briefly, peptides were labeled in DMSO at 4°C with approximately 1.3-fold molar excess of Oregon Green 488 maleimide (Life Technologies/Molecular Probes #O6034, USA) dissolved in a 1:1 solution of acetonitrile:DMSO. The reaction progress was monitored by HPLC, and labeled peptides were separated from free dye and residual unlabeled peptides using the same reversed phase conditions described above. Confirmed peptide fractions were dried under vacuum, redissolved in water, and concentrations were calculated using the absorbance of Oregon Green 488 at 491 nm (Extinction coefficient 83,000 cm^-1^ M^-1^ in 50 mM potassium phosphate, pH 9).

### Fluorescence polarization (FP)

FP experiments with IST1_NTD_ were performed in binding buffer: 25 mM Tris, pH 7.2, 300 mM NaCl, 0.1 mg/mL Bovine Serum Albumin (BSA), 0.01% Tween-20, and 1 mM Dithiothreitol (DTT) using 5–10 nM fluor-labeled ESCRT-III C-term peptides and dilutions of IST1_NTD_ and IST1_NTD_ Y165A. FP was measured using a Biotek Synergy Neo Multi-Mode plate reader (Biotek, USA) with excitation at 485 nm and detection at 535 nm. Dissociation constants were calculated by fitting the increase in FP to a 1:1 binding equation using *KaleidaGraph* (Synergy Software) as described previously (Skalicky et al., 2012). Each binding isotherm was measured at least three times independently, and mean *K*_D_ values are reported ± SD.

### Surface Plasmon Resonance

We performed Biosensor binding experiments as previously reported (Stuchell-Brereton et al., 2007). Briefly, GST-CHMP1B proteins were expressed and captured directly from BL21-Codon Plus (DE3) RIPL (Agilent, Santa Clara, CA, USA) *E. coli* cell extracts onto anti-GST antibody-derivatized CM5 sensor chips. Purified IST1_NTD_ protein (100 μM) was diluted in a 1.5-fold dilution series in binding buffer (50 mM Tris (pH 7.0), 300 mM NaCl, 1 mM DTT, supplemented with 0.01% (v/v) Tween-20 and 0.1 mg mL-1 BSA), injected in triplicate (50 μL per min, 20° C) and binding data were collected at 2 Hz during the 12–30 second association and dissociation phases. All interactions reached equilibrium rapidly and dissociated within seconds during the dissociation phase. Dissociation constants were obtained by fitting the equilibrium responses to 1:1 binding models (**Figure 4-figure supplement 1A**).

### Nuclease Protection Assay

100 μl aliquots of 50 mM Tris pH 7.0, 350 mM NaCl, 5% glycerol, 5 mM β-mercaptoethanol containing 1 μg Kpn1 linearized pUC19 dsDNA in the absence or presence of 32 μM CHMP1B and IST1_NTD_ were dialyzed overnight at room temperature against 25 mM Tris pH 8.0, 25 mM NaCl. Naked dsDNA or dsDNA:protein complexes were incubated at room temperature with either A) DNAse I (NEB #M0303S, USA) supplemented with 0.5 mM CaCl_2_ and 2.5 mM MgCl_2_ at a final concentration of 0.05 U/μg DNA or B) Micrococcal Nuclease (NEB #M0247S, USA) supplemented with 5 mM CaCl_2_ at a final concentration of 2 U/μg DNA. Reactions were quenched (5, 10, 20, or 40 mins later) by mixing with 10 mM EDTA, 10 mM EGTA, 150 mM NaCl, 200 μl Phenol/Chloroform/Isoamyl Alcohol solution. To recover the nucleic acid the following procedure was followed; The reactions were subjected to centrifugation at room temperature for 10 mins at 16,000 g, the aqueous phase recovered, 100 μl Phenol/Chloroform/Isoamyl Alcohol and the solution centrifuged at room temperature for a further 10 mins at 16,000 *g*. The resulting aqueous phase was treated with 10 μl glycogen (5 mg/mL stock), 0.1 volumes 3M NaAc (pH 5.5) and 0.7 volumes Isopropyl alcohol, followed by centrifugation at 16,000 *g* at 4° C for 30 mins. The supernatant was aspirated and the pellet vortexed in 500 μl ice cold 70% ethanol followed by centrifugation at 16,000 *g* for 10 mins. The resulting supernatant was aspirated and the pellet vortexed in 20 μl 10 mM Tris pH 7.4, 50 mM NaCl followed by incubation for 1 hour at 37° C and further vortexing. The entire 20 μl of each reaction was run in a 1% agarose gel.

### Nucleic Acid Copolymer Filament Assembly for Structure Determination

100 μl aliquots of 50 mM Tris pH 7.0, 350 mM NaCl, 5% glycerol, 5 mM β-mercaptoethanol containing 4 μM CHMP1B_4–199_, 4 μM IST1_NTD_ and the corresponding nucleic acid was dialyzed overnight at room temperature against 25 mM Tris pH 8.0 at 4 °C, 125 mM NaCl. The reactions were subjected to a slow speed spin at 2152 *g* for 10 minutes at room temperature. 75 μl of the supernatant was removed, and the pelleted filaments were resuspended with the remaining 25 μl, effectively concentrating the reaction four fold. The ssDNA nucleic acid substrate utilized was a 200 base Ultramer (IDT, USA) with a repetitive AG sequence to prevent any double-stranded nucleic acid species from forming in solution. dsDNA substrate was pUC19 plasmid DNA linearized with Kpn1 endonuclease (NEB, USA). ssRNA substrate was generated by in vitro transcription of a 1686 base sequence according to the protocol described in (Osuna et al., 2017). Excess nucleic acid was used in all copolymerization reactions for structure determination to ensure close to 100 *%* occupancy within the IST1-CHMP1B polymer for averaging purposes. All nucleic acids samples were in a 1:20 (mol:mol) ratio of one heterodimer of IST1-CHMP1B to either 20 bases (ssDNA and ssRNA) or 20 base pairs (dsDNA). Protein only assemblies were used for **Fig4-figure supplement 1B-D**. CHMP1B_1–199_ forms identical assemblies with nucleic acids as CHMP1B_4–199_ (data not shown). CHMP1B_4–199_ was chosen for structure determination as it is contains the same residues used for the binding assays throughout this study.

### Negative Stain Imaging

Nucleic acid-nucleated copolymers of IST1 and CHMP1B were prepared for TEM following established procedures (Booth et al., 2011; Grassucci et al., 2007). Imaging of negatively stained grids at the University of Utah Electron Microscopy Core Laboratory was performed as previously described (McCullough et al., 2015). Negatively stained samples imaged at the UCSF Keck Advanced Microscopy Laboratory were made with 0.75 % (w/v) uranyl formate (Structure Probe, Inc., USA) on glow-discharged carbon coated 200 Cu mesh grids (Electron Microscopy Supplies, USA). All samples were imaged on an FEI Tecnai 120 kV electron microscope equipped with a Gatan OneView CCD camera. The following reaction conditions were used for the described negative stain EM figure panels. **Figure 1A-D**, 25 mM Tris pH 8.0 and 125 mM NaCl buffer conditions with 4 μM CHMP1B_4–199_ and 4 μM IST1 FL with the corresponding nucleic acid substrate. **Figure 4 – figure supplement 1B**, 25 mM Tris pH 8.0 and 25 mM NaCl buffer conditions with 32 μM CHMP1B_4–199_ and 32 μM IST1_NTD_. **Figure 4 – figure supplement 1C**, 25 mM Tris pH 8.0 and 25 mM NaCl buffer conditions with 32 μM CHMP1B_ΔMIM_ (CHMP1B_4–178_) and 32 μM IST1_ntd_. **Figure 4 – figure supplement 1D**, 25 mM Tris pH 8.0 and 25 mM NaCl buffer conditions with 32 μM CHMP1B_4–199_ and 32 μM IST1_NTD_ Y165A.

### Vitrification Settings

Assembled nucleic acid copolymers of IST1 and CHMP1B were applied in 3.5 μL volumes to glow-discharged R1.2/1.3 Quantifoil 200 Cu mesh grids (Quantifoil, Germany) in a Mark III Vitrobot (FEI, USA). Grids were blotted with Whattman #1 filter paper (Whattman, USA) for 2–4 seconds with a 0 mm offset at 19 °C and 100 % humidity before plunging into liquid ethane. Grids were stored under liquid nitrogen until samples were imaged for structure determination.

### Electron Cryo-Microscopy Imaging

#### ssDNA Copolymer

ssDNA bound samples were imaged on a Titan Krios (FEI, USA) operating at 300 kV equipped with at K2 Summit direct electron detector (Gatan, Inc., USA) at the Berkeley Bay Area CryoEM Facility. Images were collected with a 50 μm C2 and 100 μm objective aperture with a nominal magnification of 29,000 corresponding to a 0.419 Å/pixel in super-resolution mode with *SerialEM* acquisition software (Mastronarde, 2005). To record the micrographs, 32 frames were collected at five frames per second for a 6.4 second exposure with a total electron dose of ~62 e^-^/Å^2^. 2971 micrographs were collected over a four-day session. Micrograph sums corresponding to this dataset have been deposited at EMPIAR with deposition code 223.

#### dsDNA Copolymer

dsDNA copolymer samples were imaged on a Tecniai 20 (FEI, USA) operating at 200 kV equipped with a K2 Summit direct electron detector (Gatan, Inc., USA) at the University of Utah. Images were collected with a 30 μm C2 and 100 μm objective aperture with a nominal magnification of 29,000 corresponding to a 0.617 Å/Pixel in super-resolution mode with *SerialEM* acquisition software (Mastronarde, 2005). To record the micrographs, 40 frames were collected at five frames per second for an eight second exposure with a total dose of ~53 e^-^/Å^2^.

### Data Processing and Helical Reconstruction

Dose-fractionated super-resolution raw stacks were motion corrected with *MotionCorr2* removing the first two frames and binning the micrographs by a factor of two (Zheng et al., 2017). The accumulated dose after removing the first two frames in the motion corrected sums is 51.4 e^-^/Å^2^. Non dose-fractionated micrographs were CTF corrected using *Gctf_v1.06* with the high-resolution search option from 15–4 Å (Zhang, 2016). For all subsequent data processing steps, the dose-fractionated micrographs were utilized. Filaments were manually picked with the helical picking tools within *RELION2.1* from 2980 micrographs. Overlapping particle segments were extracted into 400 pixel boxes using the extraction tools within *RELION2.1* (He and Scheres, 2017; Kimanius et al., 2016). Each particle box contained one unique turn of the helix by setting the particle overlap to 51 Å (3 Å rise with 17 subunits per turn) or a ~85% overlap. 2D classification of the 224,252 extracted particles was conducted over several rounds to remove particles that contained ice contamination and particles with intersecting filaments. The remaining 221,808 particles (98.9% of the extracted particle set) were then processed with the *RELION2.1* 3D auto-refinement procedure with a 310 pixel spherical mask. The reference model was a 60 Å low pass map generated from the protein only IST1-CHMP1B copolymer structure EMDB:6461 (McCullough et al., 2015). Helical symmetry was imposed with starting parameters of 3 Å rise and 21 ° twist and allowed to refine during the reconstruction. The consensus reconstruction from this procedure was then processed with a protein only soft mask in a 3D skip align classification (k=3) to identify the most homogeneous set of particles for refining the protein portion of the reconstruction. After multiple rounds of 3D skip align classification the class with the most homogenous high-resolution features across the distance of the volume was chosen. This portion of the dataset contained 101,990 particles (46.0 % of the original extracted particles). This class was 3D auto-refined with a soft protein only mask that was generated using relion_mask_create and the central 30% of the helix (helical_z_percentage=0.3). This refinement resulted in a 3.17 Å reconstruction following gold standard refinement procedure (Scheres and Chen, 2012). Resolutions were estimated using Fourier shell correlation (FSC), 0.143 criterion (Rosenthal and Henderson, 2003). The unfiltered half maps were post-processed within *RELION2.1* using the refinement mask and the auto-bfactor function resulting in a reconstruction filtered to 2.94 Å resolution with a temperature factor of -113 imposed on the map (Rosenthal and Henderson, 2003). The determined helical symmetry (21.1563° twist and 3.17327 Å rise) was imposed onto the asymmetric post-processed map within *relion_helix_toolbox.* Local resolution was estimated with *LocalRes* within the *RELION2.1* and had a resolution range of 2.8 to 3.2 Å (Chen et al., 2013). The density map and corresponding files for the cryoEM reconstruction have been deposited at the EMDB with depostion code EMD-9005.

### Molecular Modeling and Validation

Due to the high quality of the cryoEM map, the initial IST1-CHMP1B model was manually built in *Coot* (Emsley and Cowtan, 2004). The models were then refined with *phenix.real_space_refine* (Adams et al., 2010) using global minimization and simulated annealing. After generating initial refined models, 36 copies of IST1 and CHMP1B were generated in real space manually in *Chimera* corresponding to slightly more than two complete turns (~2.1) of the helix. Noncrystallographic symmetry (NCS) constraints were then determined based on this symmetrized model using the *find_NSC* and *apply_NSC* tools in phenix. The reference chains for the NCS operators of IST1 and CHMP1B were situated in between overlapping turns of the helices so all quaternary molecular interactions were satisfied. The symmetrized model was manually adjusted in *Coot* and further refined with *phenix.real_space_refine* using global minimization, secondary structure restraints, and local grid search. This model was manually adjusted a third and final time in *Coot* and minimized in *phenix.real_space_refine* with per-residue B-factors. Model statistics were tabulated using *Molprobity* (Chen et al., 2015) and *EMRinger* (Barad et al., 2015). **Table 1** lists all of the validation parameters for the reconstruction. Map versus atomic model FSC plots were computed using *EMAN2.1* (Ludtke, 2016) using calculated density maps from *e2pdb2mrc.py* with per-residue B-factor weighting. The final atomic model has been deposited at the PDB with deposition code 6E8G.

## ACKNOWLEDGEMENTS

We thank the following scientists at the University of Utah Health Sciences Center Core facilities for valuable assistance and advice: Mike Hanson and Scott Endicott (DNA/Peptide Synthesis Core), David Belnap (Electron Microscopy Core), Krishna Parsawar (Mass Spectrometry and Proteomics Core). Mass spectrometry equipment was obtained through NCRR Shared Instrumentation Grant # 1S10RR020883-01 and 1S10 RR025532-01A1. We thank Michael Braunfeld, Matt Harrington, David Bulkley, and Alexander Myasnikov and the UCSF Center for Advanced CryoEM and Daniel Toso and Paul Tobias of the Berkeley Bay Area CryoEM Facility. The Bay Area CryoEM consortium and the UCSF Center for Advanced CryoEM are supported in part from NIH grants S10OD020054 and 1S10OD021741 and the Howard Hughes Medical Institute. We also thank the QB3 shared cluster and NIH grant 1S10QD021596-01. The Titan X Pascal used for this research was donated by the NVIDIA corporation. This work was further supported by a Faculty Scholar grant from the Howard Hughes Medical Institute (A.F.), NIH grant 1DP2GM110772-01 (A.F.), the Sandler Family Foundation through the UCSF Program for Breakthrough Biomedical Research, the American Asthma Foundation and NIH grants P50 082545 (A.F.) and R01 GM112080 (W.I.S.). A.F. is a Chan Zuckerberg Biohub investigator.

